# A novel, evolutionarily conserved inhibitory circuit selectively regulates dentate gyrus mossy cell function

**DOI:** 10.1101/2025.07.09.663808

**Authors:** Geoffrey A. Vargish, Haley Rice, Xiaoqing Yuan, Steven Hunt, Brendan E. Hines, Anya Plotnikova, Arya Mohanty, Elisabetta Furlanis, Yating Wang, Min Dai, Brenda Leyva Garcia, Alex C. Cummins, Mark A. G. Eldridge, Bruno B. Averbeck, Kareem A. Zaghloul, Jordane Dimidschstein, Gord Fishell, Kenneth A. Pelkey, Chris J. McBain

## Abstract

The mammalian dentate gyrus contributes to mnemonic function by parsing similar events and places. The disparate activity patterns of mossy cells and granule cells is believed to enable this function yet the mechanisms that drive this circuit dynamic remain elusive. We identified a novel inhibitory interneuron subtype, characterized by VGluT3 expression, with overwhelming target selectivity for mossy cells while also revealing that CCK, PV, SOM and VIP interneurons preferentially innervate granule cells. Leveraging pharmacology and novel enhancer viruses, we find that this target-specific inhibitory innervation pattern is evolutionarily conserved in non-human primates and humans. In addition, in vivo chemogenetic manipulation of VGluT3+ interneurons selectively alters the activity and functional properties of mossy cells. These findings establish that mossy cells and granule cells have unique, evolutionarily conserved inhibitory innervation patterns and suggest selective inhibitory circuits may be necessary to maintain DG circuit dynamics and enable pattern separation across species.

## Main

The mammalian hippocampus contributes to spatial information processing and encoding of episodic memories^1–3^. Fundamental to these functions is the ability to discriminate and store perceptually similar places or events, a process that requires transforming largely overlapping input patterns of neuronal activity into discrete outputs. Anatomical, functional and theoretical evidence indicates that this computation, known as pattern separation, is performed by the dentate gyrus (DG), the primary input region of the hippocampus and the first stage in the canonical trisynaptic circuit^4–8^. The DG is comprised of two anatomically and functionally distinct classes of excitatory neurons, granule cells (GCs) and mossy cells (MCs), as well as a diverse array of inhibitory GABAergic interneurons^9–11^. Historically, pattern separation was believed to be driven by GCs, the largest cell population in the DG. GCs have sparse activity patterns, lack recurrent connectivity and exhibit prominent “expansion recoding,” outnumbering inputs from the entorhinal cortex by a factor of five; these features enable GCs to increase the dimensionality of their inputs and segregate even marginally disparate input patterns^9,12^. However, recent methodological advances that have allowed in vivo interrogation of MCs have revealed that MCs may play a critical role in pattern separation as well^13–15^.

MCs, which reside in the DG hilus, form a recurrent circuit with GCs, receiving prominent excitatory input from GCs while driving both excitation of GCs through monosynaptic input in the inner molecular layer and inhibition via strong innervation of interneurons in the DG^10,16–18^. In contrast to GCs, MCs are highly active in vivo and form multiple place fields, frequently remapping between environments^13–15^. These properties, coupled with MCs prominent innervation of DG INs, which exerts a net inhibitory effect on GCs, suggest MCs directly contribute to GC sparseness through the recruitment of disynaptic inhibition^18,19^. This indirect inhibitory feedback circuit driven by MCs could facilitate pattern separation by increasing GC signal-to-noise ratio, keeping “off-beam” GCs silent while preserving “on-beam” GC activity^20^. Consistent with this theory, ablation or inhibition of MCs impairs contextual discrimination and increases GC excitability^21,22^. Thus, at a population level, high MC activity combined with sparse GC activity presents a tractable circuit representation of DG-mediated pattern separation. We hypothesize that this circuit dynamic is generated by differential regulation of MC and GC excitability by distinct inhibitory innervation. While inhibitory inputs to GCs have been well characterized, the source of inhibition on MCs has not yet been established. Previous studies suggest that cholecystokinin (CCK)-interneurons may provide inhibitory input to MCs as morphological characterization uncovered a subset of hilar-targeting CCK interneurons and CCK immunoreactive axonal boutons were observed in apposition to MC somas, but functional evidence of CCK interneuron to MC connectivity is lacking^23,24^.

In this study we use a combination of electrophysiology, optogenetics, and in vivo 2-photon imaging to assess the organization and functional impact of inhibitory circuits in the DG, probing the source of afferent inhibitory input onto MCs. We find that inhibitory circuits in the DG exhibit unique target specificity with a novel population of vesicular glutamate transporter 3 (VGluT3) expressing interneurons predominantly innervating MCs while parvalbumin (PV), somatostatin (SOM), vasointestistinal peptide (VIP) and canonical hilar-commissural associational path (HICAP) CCK interneuron populations preferentially innervate GCs. Remarkably, whole cell patch clamp interrogation of inhibitory innervation patterns in nonhuman primate (NHP) and human tissue revealed that these inhibitory circuit motifs are evolutionarily conserved. In vivo chemogenetic manipulation of VGluT3+ interneurons led to diametric alterations in MC and GC activity while also disrupting spatial tuning properties and contextual remapping of MCs suggesting that selective inhibitory circuits play a critical role in DG circuit function across species.

## Results

### VGluT3+ interneurons are a novel hilar-targeting interneuron subtype in the DG

Given previous observations of hilar-targeting CCK+ interneurons and CCK immunoreactive axon terminals in the hilus, we first sought to examine the distribution of CCK interneuron axonal boutons in the DG that stain positive for cannabinoid receptor subtype 1 (CB1) and VGluT3, molecular markers for CCK axon terminals ^23–26^. While CB1+ axon terminals were found in the inner molecular layer, where canonical hilar-commissural projecting (HICAP) CCK interneurons innervate, and the hilus, surprisingly, VGluT3+ axon terminals were almost exclusively localized to the hilus, suggesting VGluT3 expression may be a specific marker for a novel subpopulation of hilar-targeting CCK interneurons (Fig 1A-B). As VGluT3+ serotonergic projections from the median raphe nucleus also innervate the hilus, we evaluated whether hilar VGluT3+ axonal boutons arise from local GABAergic interneurons by co-staining for VGluT3, CB1 and vesicular GABA transporter (VGAT)^27–29^. Indeed, we found a subset of hilar VGluT3+ axon terminals co-expressed both CB1 and VGAT, confirming VGluT3 is expressed in a subpopulation of CCK+ GABAergic interneurons in DG (Fig 1A-B). To probe whether hilar VGluT3+ axon terminals contact MCs, we injected VGluT3-Cre mice in contralateral DG with a retrograde adeno-associated virus (AAV2-retro) containing mCherry to selectively label MCs while staining for VGluT3 and injecting a Cre-dependent GFP virus in ipsilateral DG to selectively label DG VGluT3+ cells. Consistent with potential VGluT3+ innervation of MCs, we found VGluT3+/GFP+ boutons in apposition to virally labelled MC somas and dendrites, with VGluT3+ axon terminals preferentially forming contacts with the perisomatic region of MCs (Fig 1E).

**Figure 1.**
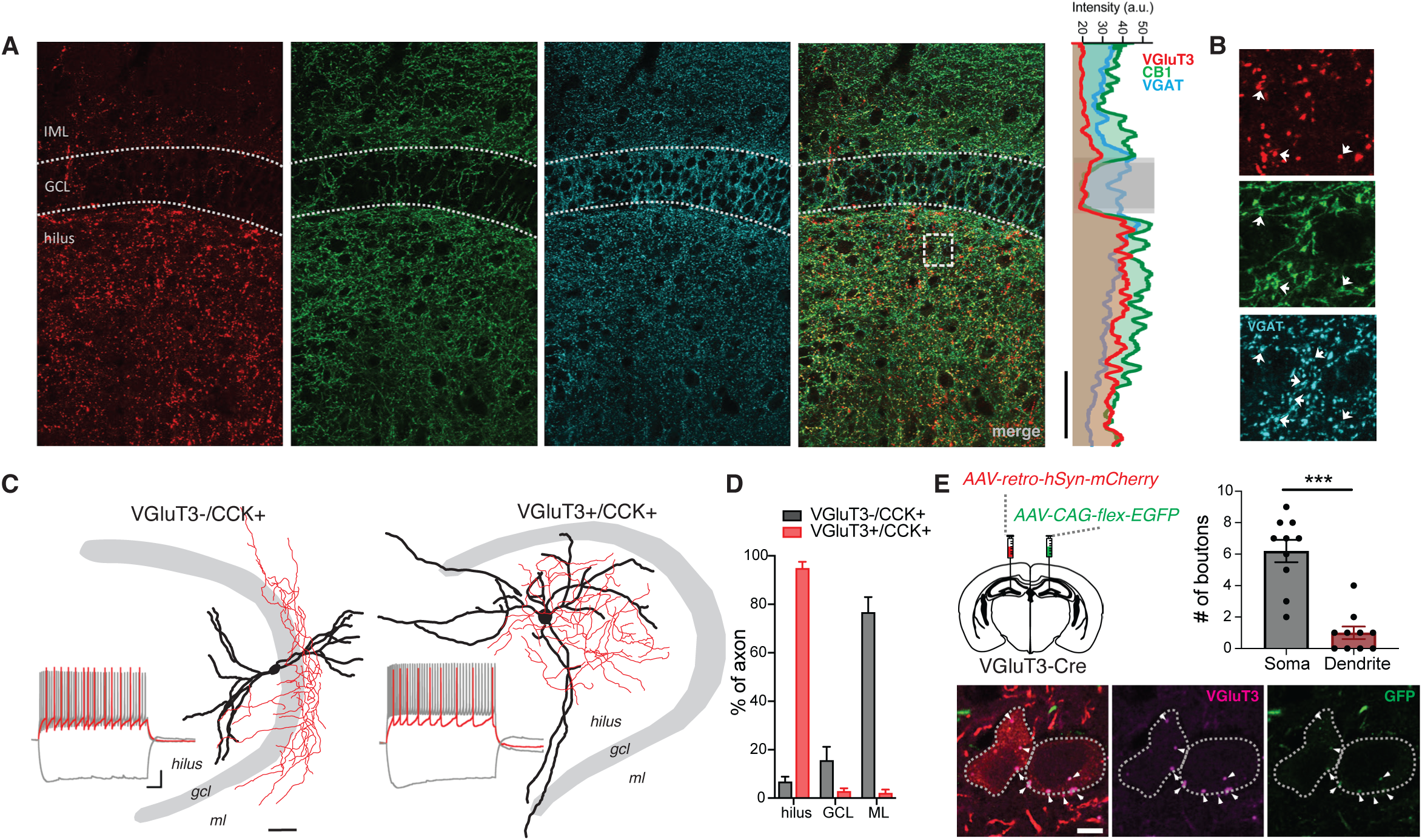
Axon terminal localization and morphology of VGluT3+ interneurons in DG. **A)** Airyscan images of fluorescent immunostaining for VGluT3 (left, red), CB1 (middle left, green) and VGAT (middle right, cyan). Profile plot (right) shows intensity of VGluT3, CB1 and VGAT signals across all layers of the DG (scale bar, 100 μm). **B)** Zoomed in images highlighting colocalization of VGluT3, CB1 and VGAT in DG hilus (scale bar, 10 μm). **C)** Posthoc morphological reconstruction of VGluT3-/CCK+ and VGluT3+/CCK+ interneurons showing axon (red) and dendrite (black) localization. Inset shows voltage response to hyperpolarizing step and spiking at threshold and at maximum firing rate. (sale bar, 50 μm). **D)** Group data depicting proportion of axon localized to hilus, GCL or ML (VGluT3-/CCK+ n = 5 cells, 2 mice; VGluT3+ n = 7 cells, 3 mice; mean ± s.e.m.). **E)** Diagram of viral injection strategy (top right). Representative images showing MCs labelled in red, VGluT3 immunostain in magenta and Cre-dependent GFP in green. Arrows show VGluT3+/GFP+ boutons in contact with MC somas (bottom). Graph shows number of axonal contacts on MC somas and dendrites (n = 10 cells, 2 mice; ***p<0.001, t-test; mean± s.e.m.).

In CA1 and CA3, VGluT3+ interneurons are morphologically and physiologically indistinguishable from the broader CCK interneuron population, comprising a portion of all anatomically distinct, classically defined CCK+ interneuron subtypes^25^. However, the unique distribution of VGluT3+ axon terminals in the DG indicates DG VGluT3+ interneurons may represent a distinct CCK+ interneuron subpopulation. To further characterize VGluT3+ interneurons and compare this cell population to other CCK+ interneurons in DG, we performed whole cell patch clamp recordings, assessing intrinsic electrophysiological properties and cell morphology. VGluT3+ interneurons were identified using VGluT3^Cre^:tdTomato^fl/fl^ mice, recording tdTomato+ cells, while VGluT3-/CCK+ cells were identified using VGluT3^Cre^:tdTomato^fl/fl^ mice crossed with either 5HT3A-GFP mice or SNCG^Flpe^ mice injected with a Flp-dependent GFP reporter virus and recording GFP+/tdTomato-cells. Corroborating our immunohistochemical bouton localization, posthoc morphological reconstruction of biocytin-filled cells revealed that VGluT3+ interneurons have axon clouds almost entirely restricted to the hilus (94.9%) while VGluT3-/CCK+ interneurons predominantly innervated the inner molecular layer (IML) (76.7%), with only 6.8% of axon distributed in the hilus (Fig 1C-D). In addition, while VGluT3+ and VGluT3-/CCK+ interneurons had similar resting membrane potential, action potential kinetics, and adaption ratios, VGluT3+ interneurons exhibited lower input resistance and a faster membrane time constant when compared to the VGluT3-/CCK+ population (Supp. Fig 1K-T). Together these findings indicate that VGluT3+ interneurons comprise a novel, morpho-physiologically distinct subpopulation of interneurons in the DG that selectively target the hilus, representing a departure from CA1 and CA3 local microcircuitry^25^.

### VGluT3+ interneurons selectively innervate MCs

The presence of VGluT3+ terminals surrounding MCs as well as the localization of VGluT3+ interneuron axons almost exclusively in the hilus indicates that VGluT3+ interneurons may exhibit target selectivity for MCs. To determine the postsynaptic target specificity of DG VGluT3+ interneurons, we performed paired whole cell patch clamp recordings of a presynaptic VGluT3+ interneuron and postsynaptic MC or GC, driving action potentials in the presynaptic neurons while assessing unitary inhibitory postsynaptic currents (IPSCs) in postsynaptic cells. The connectivity rate of VGluT3+ interneuron to MC pairs was a remarkable 73.3% compared to only 25% for VGluT3+ interneuron-GC pairs (Fig 2A-D). Further, the amplitude of VGluT3-mediated uIPSCs was significantly larger in MCs when compared to GCs (Fig 2E). The large amplitude VGluT3-mediated uIPSCs recorded from MCs (569pA ± 157pA) is consistent with our anatomical observations that VGluT3+ interneuron axon terminals innervate MCs perisomatically (Fig 2E). In contrast to VGluT3+ interneurons, VGluT3-/CCK+ interneurons lacked connectivity with MCs but had a 27.3% connectivity rate with GCs suggesting MC target selectivity is a specific feature of VGluT3+ interneurons (Fig 2B-D).

**Figure 2.**
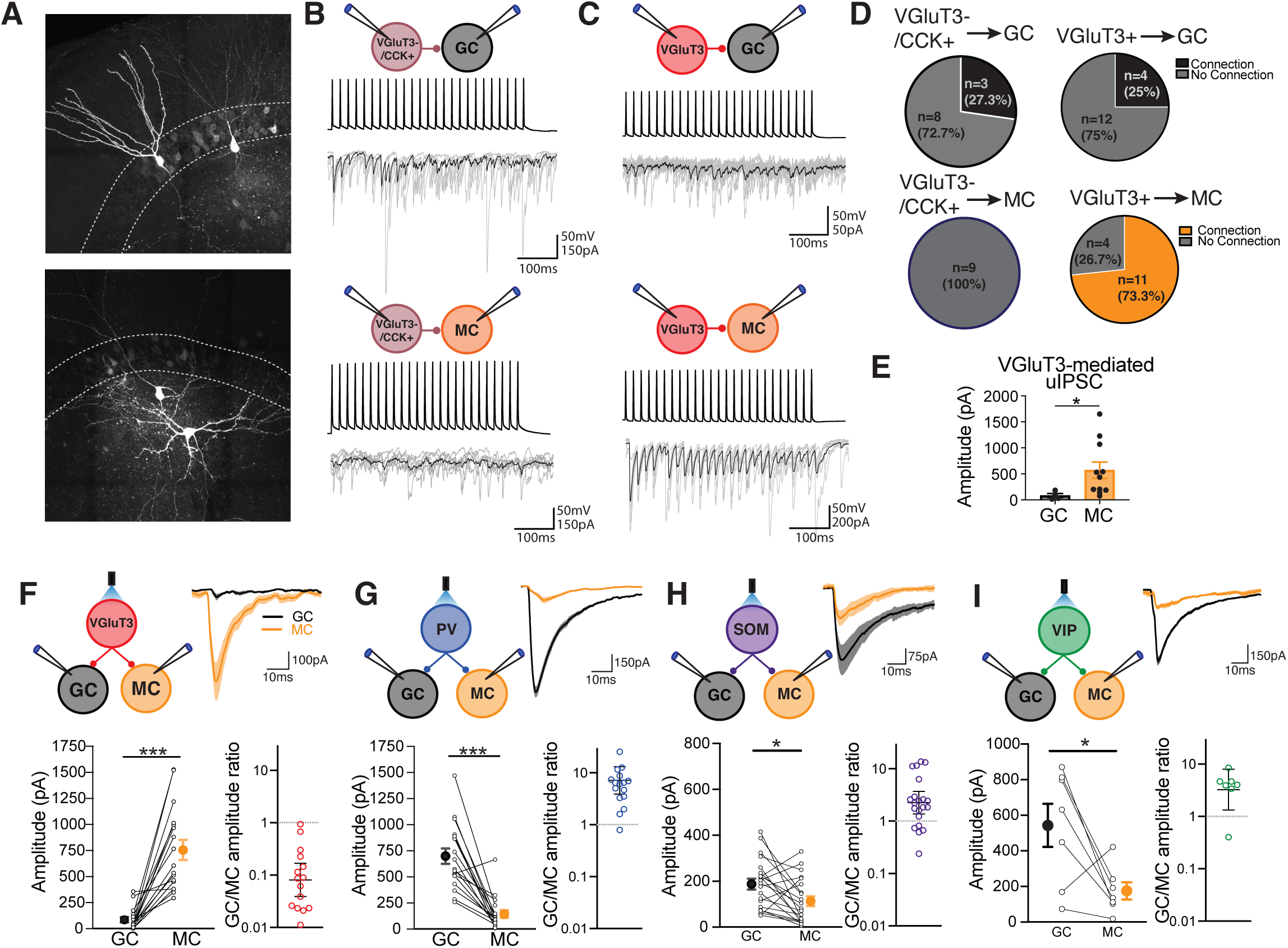
Target selective inhibitory connectivity in the DG. **A)** Representative images of biocytin filled VGluT3+ interneuron-GC (top) and VGluT3+ interneuron-MC (bottom) pairs. **B-C)** Diagram of recording setup and representative traces of current clamp recordings from presynaptic interneuron and voltage clamp recordings from postsynaptic neuron for VGluT3-/CCK+ (B) and VGluT3+ (C) interneuron paired recordings with postsynaptic GCs (top) and MCs (bottom) **D)** Pie charts showing percent connectivity for the different paired recording configurations highlighted in B and C. **E)** Group data showing uIPSC amplitude for VGluT3+ interneuron-GC and VGluT3+ interneuron-MC pairs (VGluT3-GC, n = 4 pairs, 4 mice; VGluT3-MC, n = 11 pairs, 6 mice; p=0.0103, Mann-Whitney U test; mean± s.e.m.). **F-I)** Diagram of recording setup (top left) and representative leIPSC traces (top right) recorded in GCs and MCs after photostimulation of VGluT3 (F), PV (G), SOM (H) and VIP (I) interneurons. Bottom left graphs show group data and individual values for leIPSCs simultaneously recording in MCs and GCs following photostimulation of the different interneuron subtypes, mean± s.e.m.. Bottom right graphs show ratio of leIPSC amplitude in GCs compared to MCs depicting geometric mean±s.e.m. and individual values. (VGluT3, n=18 pairs, 4 mice; PV, n=18 pairs, 4 mice; SOM, n=21 pairs, 4 mice; VIP, n=7 pairs, 2 mice;***p<0.001,*p<0.05, paired t-test)

CCK+ interneurons in DG and CA1/CA3, including the VGluT3+ population in CA1/CA3, are characterized by distinct synaptic properties including CB1-mediated suppression of presynaptic release, asynchronous neurotransmitter release, and highly variable uIPSCs, resulting from low temporal fidelity and a high proportion of synaptic failures^25,30–33^. We investigated whether the unique morpho-physiology and target specificity of DG VGluT3+ interneurons is also associated with differences in synaptic properties when compared to the broader CCK+ population in the hippocampus. Consistent with other CCK+ subpopulations, VGluT3+ interneurons in DG exhibited asynchronous release during trains of action potentials and prominent CB1-mediated suppression of neurotransmitter release in response to both endogenous cannabinoids, triggered through depolarization-induced suppression of inhibition (DSI), and an exogenous CB1 agonist, WIN-55,212-2(Supp Fig 1A-J)^25,30^. Surprisingly, though, we found that VGluT3+ interneurons in the DG had higher release probability, as indicated by lower failure rate and uIPSC coefficient of variation (CV), and lower synaptic latency with less synaptic jitter when compared to other populations of CCK+ interneurons in CA1, CA3 and DG (Supp Fig 1G&H)^31,33^. These synaptic properties may confer a different functional specialization for VGluT3+ interneurons in the DG circuit.

Given the target selectivity observed in both VGluT3+ and VGluT3-/CCK+ interneuron subpopulations, we next asked whether other interneuron subtypes in the DG also exhibit selectivity for either MCs or GCs. Using Cre driver lines specific for VGluT3, PV, SOM and VIP interneurons to express channelrhodopsin (ChR2) in these neuronal populations, we investigated the strength of inhibitory input from each interneuron subtype onto MCs and GCs. We performed dual patch clamp recordings of MCs and GCs while delivering widefield photostimulation to directly compare light-evoked IPSCs in these cells (Fig 2F-I). In line with data from paired recordings, the amplitude of light-evoked IPSCs mediated by VGluT3+ interneurons was significantly larger in MCs when compared to GCs (Fig 2F). Importantly, this difference was not a result of differential recruitment of VGluT3+ inputs onto MCs and GCs as input/output curves of light-evoked IPSCs revealed significant differences in IPSC amplitude across all light intensities (Supp Fig 2A-C). PV, SOM and VIP interneurons exhibited the opposite pattern of connectivity. Light-evoked IPSCs mediated by these interneuron subtypes were biased toward GCs displaying 7.17, 2.25 and 3.25-fold greater amplitude in GCs when compared to MCs for PV, SOM and VIP interneurons, respectively (Fig 2G-I). These findings establish that the organization of inhibitory inputs onto MCs and GCs is unique, with VGluT3+ interneurons providing the dominant source of inhibition onto MCs while VGluT3-/CCK+, PV, SOM and VIP interneuron subpopulations primarily innervate GCs.

### VGluT3+ interneurons are recruited in feedforward and feedback circuits

Our connectivity analysis revealed VGluT3+ interneurons as the principal source of inhibition onto MCs, yet it remains unclear how VGluT3+ cells are engaged in the DG circuit to ultimately shape MC excitability. Post-hoc reconstructions of VGluT3+ interneuron morphology indicates VGluT3+ interneurons have dendritic arbors that span all layers of the DG, suggesting these cells can be recruited in both the feedforward circuit, via entorhinal cortex (EC) afferents in the molecular layer or mossy fiber input from GCs in the hilus, and the feedback circuit through innervation by MCs (Fig 1C). To first assess whether VGluT3+ interneurons are innervated by MCs, we injected VGluT3-Cre:Ai14 mice with a retrograde virus containing ChR2 in contralateral DG and examined light-evoked EPSCs (leEPSCs) simultaneously in GCs and tdTom+ VGluT3+ interneurons (Fig 3A-C). Wide-field photostimulation of ipsilateral DG elicited monosynaptic leEPSCs in GCs, as expected based on prior studies^10^, and in VGluT3+ interneurons (Fig 3C-D). leEPSC amplitude was significantly larger in VGluT3+ interneurons when compared to GCs, with leEPSCs recorded in VGluT3+ interneurons 4.43-fold greater than those in GCs, indicating MCs can recruit VGluT3+ interneurons in the feedback circuit through potent monosynaptic excitation (Fig 3C-D).

**Figure 3.**
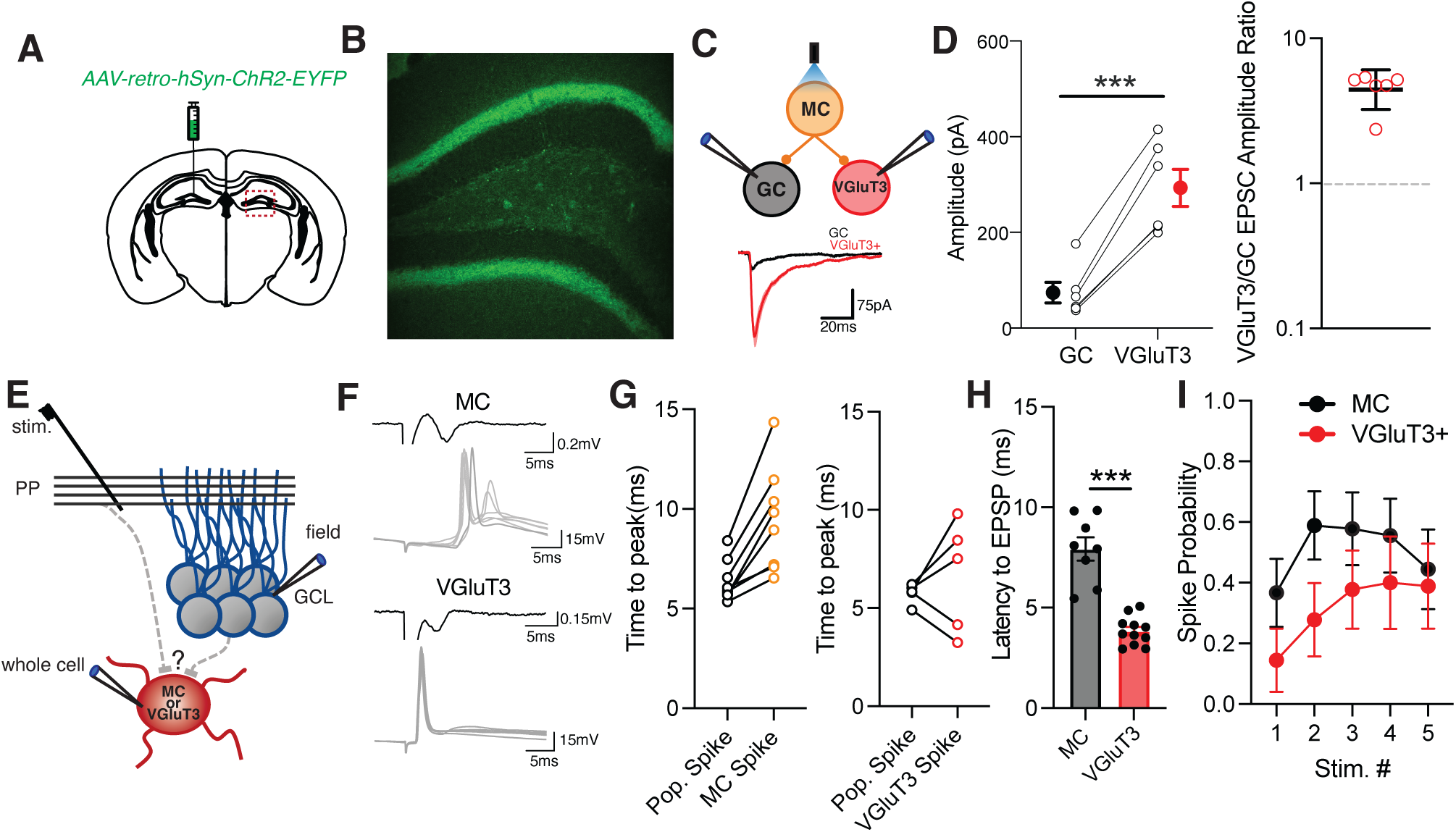
VGluT3+ interneuron engagement in feedforward and feedback circuits. **A)** Diagram of viral injection for retrograde infection of MCs. **B)** Image of contralateral DG showing infection of MCs with ChR2-YFP virus. **C)** Diagram of recording setup (top) and representative images of leEPSCs (bottom) recording from VGluT3+ interneurons and GCs after MC photostimulation. **D)** Graphs showing group data and individual values for paired recordings of leEPSCs in VGluT3+ interneurons and GCs during MC photostimulation, mean± s.e.m., (left) as well as ratio of leEPSC amplitude in VGluT3+ interneurons compared to GCs for each pair, geometric mean± s.e.m.(n=6 pairs, 2 mice; ***p<0.001, paired t-test). **E)** Schematic of recording setup for feedforward pathway experiments. **F)** Representative traces of evoked field population spike recorded in GCL (black) and whole cell evoked action potentials (gray) recorded in MCs (top) and VGluT3+ interneurons (bottom). **G)** Graphs showing latency from stimulation to evoked pop spike and whole cell action potential for MC (left) and VGluT3+ interneuron (right) recordings (MC, n=8 cells, 3 mice; VGluT3, n=5 cells, 5 mice). **H)** Group data showing latency to evoked EPSP onset in MCs and VGluT3+ interneurons (MC, n=8 cells, 3 mice; VGluT3, n=11 cells, 5 mice; ***p<0.001 t-test; mean± s.e.m.). **I)** Graph showing spike probability in MC and VGluT3+-interneurons for each stimulation in a 5-pulse train at 10Hz. mean± s.e.m.

Next, we investigated whether VGluT3+ interneurons participate in feedforward inhibition, examining VGluT3+ interneuron innervation by perforant path EC afferents and GC mossy fibers. To probe these pathways, we performed whole cell current clamp recordings of VGluT3+ interneurons, or MCs, with simultaneous field potential recordings in the GCL, and measured the timing of spikes in the whole cell recording relative to the population spike in the GCL evoked by electrical stimulation in the middle molecular layer (MML) (Fig 3E). Action potentials arriving before the GCL pop. spike indicate direct EC input while action potentials with a longer latency than the pop spike points to recruitment by the GC population. In line with observations that MCs rarely send dendrites to the ML, we found that action potentials in MCs reliably had a longer latency than the GCL pop spike (Fig 3F-G)^10^. Recordings of VGluT3+ interneurons, however, revealed that 2/5 VGluT3+ interneurons had spikes with shorter latency than the GCL pop. spike and the remaining 3/5 cells had latencies longer than the pop spike (Fig 3F-G). While this suggests there are VGluT3+ interneuron subpopulations with different afferent innervation, examination of evoked excitatory postsynaptic potential (EPSP) latency in the absence of spiking uncovered that VGluT3+ interneurons had a shorter EPSP latency when compared to MCs (Fig 3H). This indicates VGluT3+ interneurons are innervated by EC afferents but that excitation-spike coupling in VGluT3+ interneurons may be weak. Indeed, we found that spike probability in VGluT3+ interneurons in response to MML stimulation that reliably drives a GCL pop spike is low, starting at 14.4% and increasing to 38.8% across a 5-pulse train (Fig 3I). Altogether these findings demonstrate that VGluT3+ interneurons can be recruited in both perforant path and mossy fiber feedforward pathways as well as in the feedback circuit.

### DG inhibitory connectivity patterns are conserved in nonhuman primates and humans

To evaluate the translational relevance of the DG inhibitory innervation patterns we observed in rodents, we investigated inhibitory connectivity in acute DG slices from rhesus macaques and resected human tissue from pharmacoresistant epilepsy patients. Fluorescent in situ hybridization confirmed the presence of *Slc17a8* (VGluT3) expressing cells in macaque DG and showed colocalization of *Slc17a8* and *Cnr1* in the hilus (Fig 4A). To probe VGluT3+ interneuron input onto MCs and GCs, we recorded monosynaptically evoked IPSCs in each cell type and assessed the proportion of cannabinoid sensitive input using DSI (Fig 4B&D). In both macaques and humans, eIPSCs were significantly more cannabinoid sensitive in MCs when compared to GCs, consistent with VGluT3+ interneuron selective innervation of MCs (Fig 4 B-E). Notably, the magnitude of DSI-mediated eIPSC suppression in macaque and human MCs (41.4±4.4% and 37.7±1.5%) was similar to the suppression observed in rodent VGluT3+ interneuron-MC pairs (47.7±9.8%) suggesting predominant VGluT3+ interneuron innervation of MCs is conserved across species. To further probe VGluT3+ interneuron target specificity in macaques we injected adult macaques with *AAV-PHP.eB-mDlx-ChR2-mCherry* in hippocampus to express ChR2 in interneurons (Fig 4F)^34^. In line with eIPSC findings, Dlx leIPSCs recorded in MCs exhibited more sensitivity to the synthetic CB1 agonist, WIN55,212-2, than leIPSCs recorded in GCs (Fig 4G-H).

**Figure 4.**
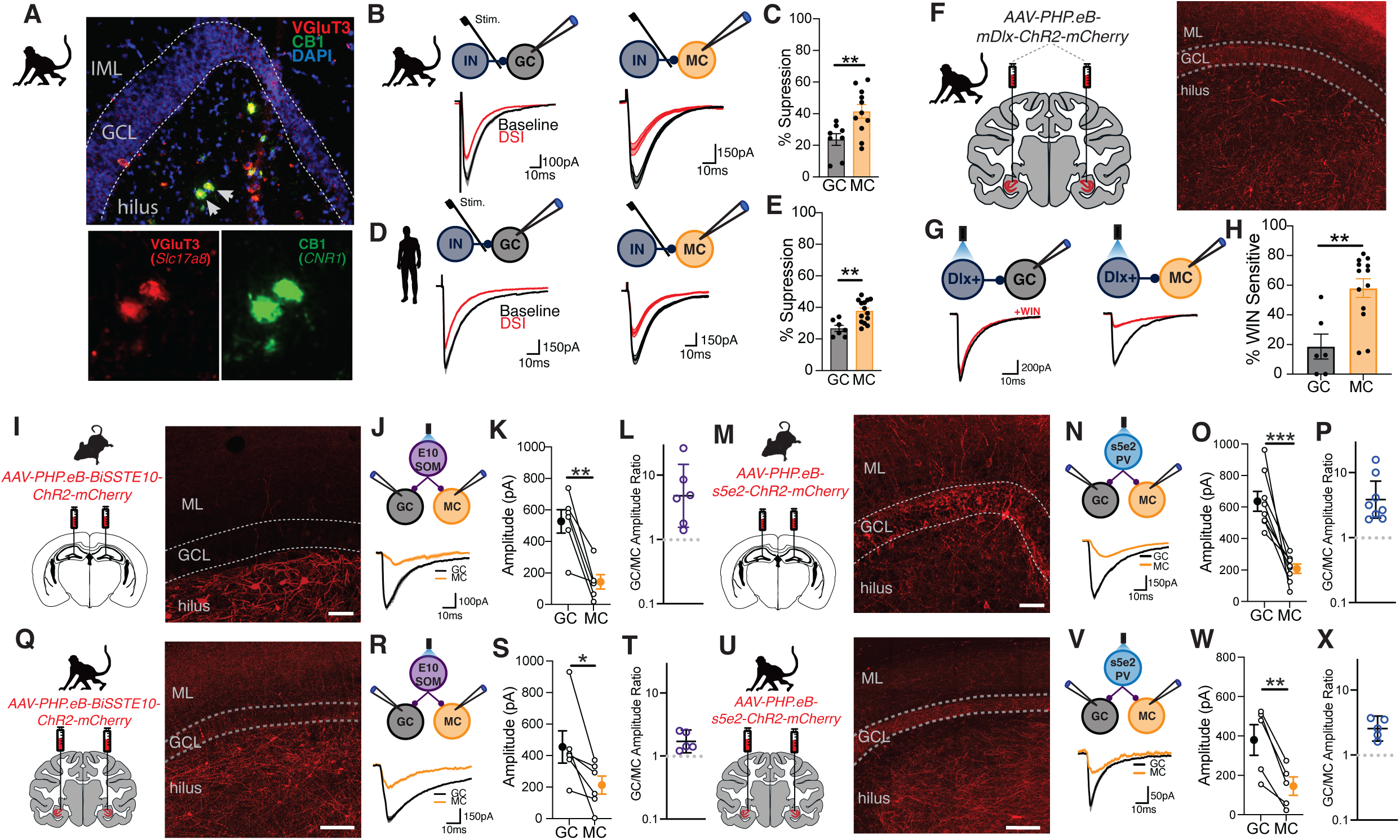
Evolutionary conservation of DG inhibitory circuit connectivity. **A)** Images of RNAScope fluorescence in situ hybridization for Slc17a8 (VGluT3) and Cnr1 (CB1) in macaque DG. Inset images show cells highlighted by arrows. **B)** Diagram of recording setup for evoked IPSC recordings from GCs and MCs in macaque DG (top) and representative traces during baseline and after DSI (bottom). **C)** Group data showing percent suppression of eIPSC after induction of DSI in GCs and MCs in macaques (GC, n=8 cells, 2 animals, MC, n=11 cells, 2 animals; **p<0.01 t-test; mean± s.e.m.). **D)** Same as in B for human tissue surgically resected from pharmacoresistant epilepsy patients. **E)** Group data showing percent suppression of eIPSC after induction of DSI in GCs and MCs in human tissue (GC, n=7 cells, 3 patients; MC, n=14 cells, 3 patients; **p<0.01, t-test; mean± s.e.m.). **F)** Diagram of bilateral viral injections of Dlx virus in macaque hippocampus and image Dlx viral expression in macaque DG. **G)** Schematic of setup for recording Dlx-mediated leIPSCs in GCs (left)and MCs (right) and representative traces of leIPSCs during baseline and after bath application of WIN55,212-2 (bottom). **H)** Graph showing percent suppression of leIPSCs in GCs and MCs after bath application of WIN55,212-2 (GC, n=6 cells, 2 animals; MC, n=13 cells, 2 animals; **p<0.01 t-test; mean± s.e.m.). **I)** Diagram of BiSSTE10-ChR2-mCherry injection in rodent vDG (left) and image of BiSSTE10-ChR2-mCherry expression in mouse DG (right). **J)** Schematic of recording setup for simultaneous GC and MC recording during photostimulation of BiSSTE10+ cells (top). Representative traces of BiSSTE10-mediated leIPSCs in GCs and MCs (bottom). **K-L)** Graphs showing BiSSTE10 leIPSC amplitude, mean± s.e.m., (K) and amplitude ratio, geometric mean±s.e.m. (L) for leIPSCs recorded in rodent GCs and MCs (n=6 pairs, 2 mice; **p,0.01, paired t-test). **M)** Same as in I but for s5e2-ChR2-mCherry injections. **J)** Recording setup and representative traces for s5e2 recordings in GCs and MCs. **O-P)** Graphs depicting s5e2 leIPSC amplitude, mean± s.e.m., and ratio of s5e2 leIPSC amplitude, geometric mean± s.e.m., in rodent GCs and MCs (n=8 pairs, 2 mice; ***p<0.001 paired t-test). **Q)** Diagram of BiSSTE10-ChR2-mCherry injections in macaque hippocampus and images of BiSSTE10 virus expression in macaque DG. **R)** Recording setup and representative traces for simultaneous GC and MCs recordings during BiSSTE10 photostimulation in macaque DG. **S-T)** Graphs showing BiSSTE10 leIPSC amplitude, mean± s.e.m., and amplitude ratio, geometric mean± s.e.m. in macaque GCs and MCs (n=5 pairs, 1 animal; *p<0.05 paired t-test). **U-V)** Same as in Q but for s5e2-ChR2-mCherry virus. **W-X)** Group data and individual values for s5e2 leIPSC amplitude, mean± s.e.m., and amplitude ratio, geometric mean± s.e.m., in macaque GCs and MCs (n=5 pairs, 1 animal; **p<0.01 paired t-test).

The recent development of novel enhancer viruses that target distinct interneuron subtypes has enabled detailed circuit dissection across species^34–36^. Using enhancer viruses targeting PV, S5E2, and SOM, BiSSTE10, interneurons to drive ChR2 expression, we investigated the target selectivity of these interneuron subtypes in macaques. Confirming the specificity of these viruses, both S5E2 and BiSSTE10 leIPSCs were blocked by bath application of bicuculline and were insensitive cannabinoids - either DSI or WIN55,212-2 (Supp. Fig 3). In addition, simultaneous MC and GC recordings in rodents injected with *AAV-PHP.eB-S5E2-ChR2-mCherry* or *AAV-PHP.eB-BiSSTE10-ChR2-mCherry* demonstrated that both S5E2 and BiSSTE10 input was stronger in GCs when compared MCs, consistent with the PV and SOM target selectivity observed using PV^Cre^ and SOM^Cre^ mice (Fig 4I-P). Intrahippocampal injection of S5E2 and BiSSTE10 ChR2 viruses in macaques revealed that S5E2 and BiSSTE10 populations exhibit the same bias towards GCs in macaques (Fig 4Q-X). S5E2 and BiSSTE10 leIPSC amplitude was 2.55- and 1.7-fold larger in GCs when directly compared to MCs, respectively (Fig 4T&X). Overall, these findings establish the evolutionary conservation of DG inhibitory circuit organization from rodents to nonhuman primates and humans with CB1+ interneurons principally innervating MCs, consistent with selective VGluT3+ interneuron innervation, while PV and SOM interneurons preferentially innervate GCs in all three species.

### Chemogenetic activation of VGluT3+ interneurons alters DG activity dynamics

While several recent studies have characterized the activity patterns, spatial representations and behavioral recruitment of MCs and GCs in vivo, understanding of the contribution of distinct interneuron subpopulations to functional properties of MCs and GCs remains limited^13–15,37,38^. Our findings that VGluT3+ interneurons exhibit target selectivity for MCs and serve as the dominant source of inhibition on MCs suggests VGluT3+ interneurons may play critical role in regulating MC function. We sought to evaluate how VGluT3+ interneurons influence in vivo activity patterns and functional properties of MCs using a combination of in vivo 2-photon imaging and chemogenetics in awake, head-fixed mice immersed in a virtual reality environment. VGluT3^Cre^ mice were injected with a CaMKII-GCaMP6f/8f AAV, to enable Ca^2+^ imaging of MCs and GCs, and a Cre-dependent AAV containing the chemogenetic activator, hM3Dq, which can reversibly activate VGluT3+ interneurons with administration of DREADD agonist 21 (C21) (Figure 5E). Following implantation of an imaging cannula allowing optical access to the DG, mice were trained to navigate a 3 m linear virtual environment while receiving sucrose rewards at specific locations along the linear track (Figure 5A-D).

**Figure 5.**
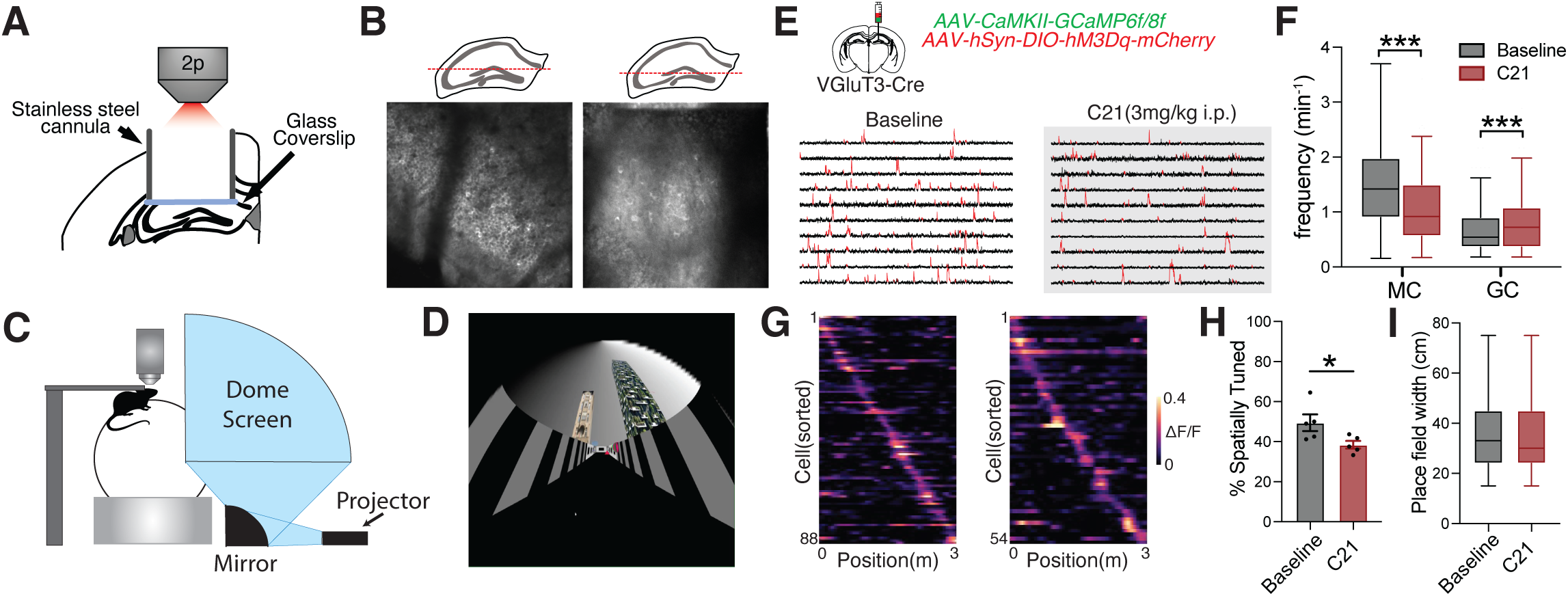
Assessing chemogenetic activation of VGluT3+ interneurons in vivo. **A)** Diagram of imaging cannula for in vivo 2-photon imaging of DG. **B)** Diagram of focal plane for GCL (left) and hilar (left) imaging (top). Time-averaged 2-photon images of GCL and hilus (bottom). **C)** Schematic of imaging setup with virtual reality. The mouse is head-fixed running on top of a floating ball while the virtual environment is projected on a dome that surrounds the mouse. **D).** Image of familiar virtual reality environment. **E)** Diagram of viral injections for chemogenetic activation of VGluT3+ cells and GCaMP imaging of MCs/GCs (top). Representative fluorescence traces from MCs during baseline (left) and after system injection of C21 (right) – significant transients are highlighted in red. **F)** Group data on Ca^2+^ transient frequency during baseline and C21 sessions for MCs and GCs (MCs: baseline n=274 cells, 7 mice, C21 n=257 cells, 7 mice; GCs: baseline n=484 cells, 3 mice, C21 n=412 cells, 3 mice; ***p<0.001 Mann-Whitney U test). **G)** Spatial tuning heatmaps for spatially tuned mossy cells in baseline and C21 sessions. Cells are sorted by the location of the peak of their spatial tuning curves. **H)** Group data showing proportion of spatially tuned cells in each mouse during baseline and C21 sessions (n=5 mice; *p<0.05 t-test, mean± s.e.m.). **I)** Group data showing MC place field width during baseline and C21 sessions (baseline, n=88 cells, 5 mice; C21, n=54 cells, 5 mice; mean± s.e.m.).

First examining overall activity patterns following VGluT3+ interneuron activation, we found that systemic administration of C21 (3 mg/kg) significantly reduced MC Ca^2+^ transient frequency and concomitantly increased GC transient frequency when compared to baseline imaging sessions (Fig 5E-F). This divergent effect on MC and GC activity confirms the VGluT3+ interneuron target specificity we observed through in vitro circuit dissection. To determine whether VGluT3+ interneurons contribute to spatial representations in MCs, we also evaluated MC spatial tuning properties after VGluT3+ activation. Chemogenetic activation of VGluT3+ interneurons reduced the fraction of MCs that exhibited spatial tuning but preserved place field width (Fig 5G-H). VGluT3+ interneuron activation also disrupted the relationship between MC activity and velocity, reducing activity-velocity correlations and altering the temporal association of MC activity to changes in velocity, with VGluT3+ interneuron activation leading to looser temporal coupling (Supp. Fig 4D-E). Together these findings demonstrate that VGluT3+ interneurons selectively regulate MC activity, spatial representations and state-dependent recruitment as well as indirectly influence GC activity patterns in vivo.

### Silencing VGluT3+ interneurons alters MC remapping

Both GCs and MCs “remap” their spatial activity profiles in response to changes in the environment, a property thought to support pattern separation^7,13,14,39^. Manipulation of PV and SOM interneurons alters remapping in GCs suggesting interneurons may contribute to this function^40^. To investigate whether VGluT3+ interneurons influence remapping of MC activity between contexts, we chemogenetically silenced VGluT3+ interneurons and monitored MC activity as mice navigated a familiar virtual environment, introduced during training sessions, and a novel virtual environment, with distinct visual cues and a different reward location (Fig 6A-B). In line with the effects of VGluT3+ interneuron activation, chemogenetic silencing of VGluT3+ interneurons with systemic administration of C21 (3 mg/kg) significantly increased MC transient frequency while reducing transient frequency in GCs when compared to baseline imaging sessions (Fig 6C). Further, silencing VGluT3+ interneurons increased the proportion of MCs expressing place fields in the familiar or novel environments but had no effect on place field width in either environment or on correlations between MC activity and behavioral state (Supp Fig 5B-E).

**Figure 6.**
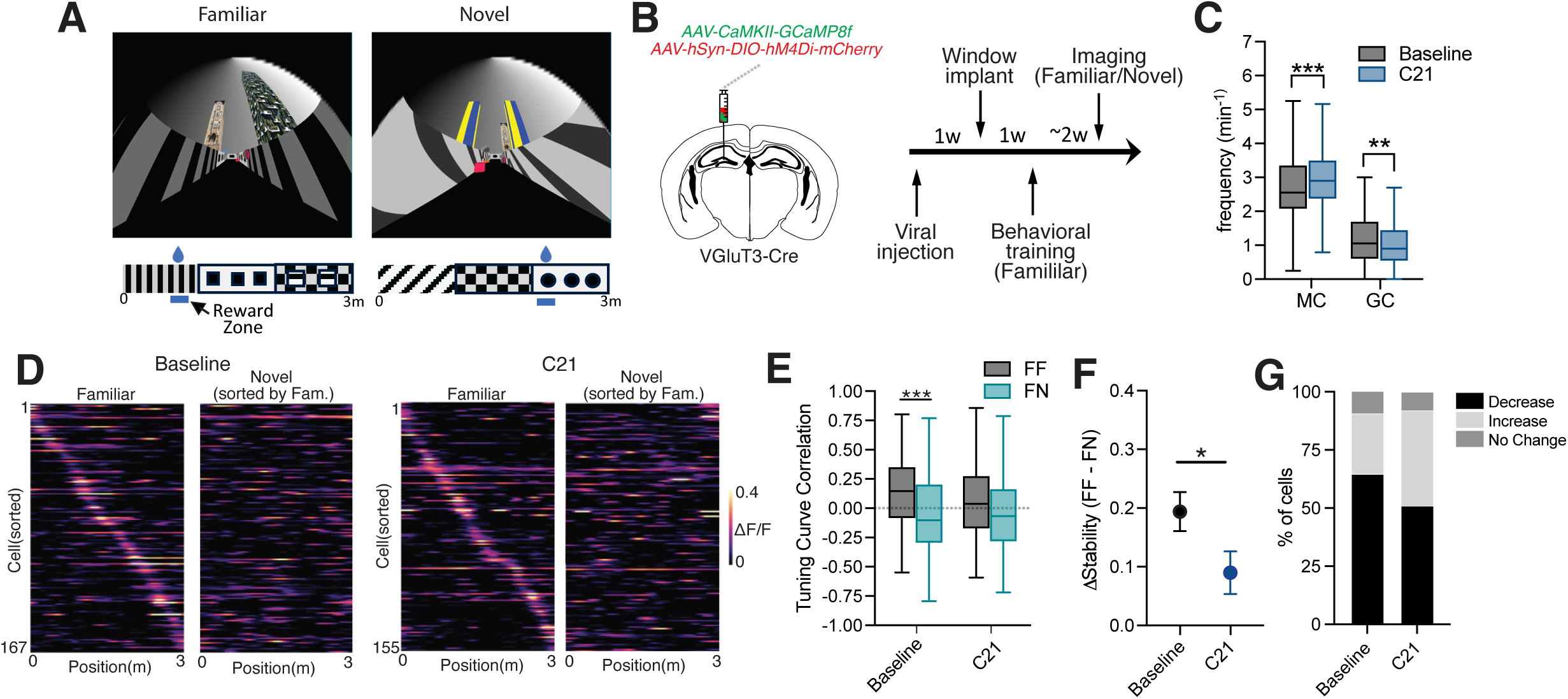
Chemogenetic inhibition of VGluT3+ interneurons alters contextual remapping. **A)** Images of familiar (left) and novel (right) virtual environments (top). Schematic of the layout and reward location for familiar and novel linear virtual environments (bottom). **B)** Schematic of viral injections for chemogenetic inhibition of VgluT3+ interneurons and imaging of MCs/GCs (left). Experimental timeline for chemogenetic inhibition during contextual switching paradigm (right). **C)** Group data showing Ca^2+^ transient frequency during baseline and C21 sessions for MCs and GCs (MCs: baseline, n=312 cells, 5 mice, C21 n=271 cells, 5 mice; GCs: baseline n=407 cells, 2 mice, C21 n=550 cells, 2 mice; ***p<0.001, **p<0.01 Mann-Whitney U test). **D)** Spatial heatmaps of MCs with a significant place field in either the familiar or novel context during baseline and C21 sessions. Cells were sorted by the peak of their spatial tuning curve in the familiar context. **E)** Group data showing tuning curve correlations for within context (FF) and between context (FN) comparisons during baseline and C21 sessions (baseline n=167 cells, 3 mice; C21 n=155 cells, 3 mice; ***p<0.001 Kruskal-Wallis test). **F)** Graph showing the average magnitude of difference in correlation (Δ stability) between FF and FN comparisons for baseline and C21 sessions (baseline n=167 cells, 3 mice; C21 n=155 cells, 3 mice; *p<0.05 t-test; mean± s.e.m.). **G)** Plot showing proportion of MCs baseline and C21 conditions that exhibit a decrease, increase or no change in tuning curve correlation from FF to FN (baseline n=167 cells, 3 mice; C21 n=155 cells, 3 mice).

To assess MC remapping, for each cell with a place field in at least one of the environments we calculated tuning curve correlations within the same virtual environment (FF, laps split in half) and between environments (FN, familiar vs novel). While FF correlations were significantly greater than FN correlations during baseline sessions, there was no difference between FF and FN correlations after silencing VGluT3+ interneurons (Fig 6 E). In addition, examining the magnitude of the change in spatial tuning stability (Δ stability) by calculating the difference between FF and FN correlations for each cell, we found that silencing VGluT3+ interneurons lead to a lower Δ stability when compared to baseline (Fig 6 F). During baseline sessions 64.7% of MCs had place fields that were more stable in FF comparisons than in FN and only 25.7% exhibited lower stability in FF than FN (9.6% had no change). In contrast, silencing VGluT3+ interneurons resulted in 50.9% of MCs displaying greater tuning curve stability in FF comparisons while 41.3% had lower stability in FF than FN (7.8% no change). These findings indicate that silencing VGluT3+ interneurons disrupts MC remapping between contexts and suggest VGluT3+ interneurons may contribute to contextual discrimination in the DG circuit.

## Discussion

Our study uncovered a novel GABAergic interneuron subtype in the DG, characterized by expression of VGluT3, with postsynaptic target specificity for MCs. In vitro circuit dissection of inhibitory inputs to MCs and GCs revealed that other DG interneuron subpopulations, including PV, SOM, VIP and CCK+/VGluT3-subtypes, also exhibit postsynaptic target selectivity, preferentially innervating GCs over MCs. This differential pattern of inhibitory innervation onto MCs and GCs explains, in part, the disparate activity patterns observed in MCs and GCs in vivo as well as supports evidence suggesting these neuronal populations are independently modulated despite forming a reciprocal excitatory circuit^13–15^. Remarkably, experiments in nonhuman primate and human tissue, leveraging CB1 expression in VGluT3+ interneurons and novel enhancer viruses selective for interneuron subpopulations, revealed that the organization of inhibitory inputs onto MCs and GCs is evolutionarily conserved. Combining chemogenetics with in vivo 2-photon imaging of Ca^2+^ transients in awake, head-fixed mice immersed in a virtual reality environment we also found that VGluT3+ interneuron manipulation directly influences MC activity patterns while leading to indirect and opposing changes in GC activity. Further, chemogenetic activation and silencing of VGluT3+ interneurons alters spatial tuning of MCs and the degree of remapping between contexts. These findings indicate that VGluT3+ interneurons contribute to the regulation of MC function and more broadly suggest that unique inhibitory innervation patterns may play a critical role in governing DG circuit function and context discrimination across species.

The unique morphology, physiology and connectivity of VGluT3+ interneurons indicates that this cell population represents a novel interneuron subtype, distinct from the VGluT3+ interneurons previously described in other hippocampal subfields. Previous studies in CA1 and CA3 have shown that VGluT3 is expressed in a subset of all canonical morphological subtypes of CCK+ interneurons and that VGluT3+ interneurons in these regions are physiologically and synaptically analogous to the VGluT3-/CCK+ population^25,26,30,41^. In CA1 and CA3 these cells are characterized by presynaptic modulation through CB1 receptors, prominent asynchronous neurotransmitter release, and highly variable synaptic release properties^25,31–33^. These features are thought to serve a gain modulation role that “fine-tunes” the hippocampal network complimenting the temporally precise feedforward input of PV interneurons that drives network synchrony^42–44^. In contrast, while VGluT3+ interneurons in DG exhibited presynaptic CB1 modulation and asynchronous release, they had higher release probability and improved temporal fidelity, as indicated by shorter synaptic latency with less jitter, when compared to other CCK+ interneuron populations (including the VGluT3+ population)^25,31–33^. These synaptic differences may reflect the minimal PV interneuron input observed in MCs. As VGluT3+ interneurons are the primary source of inhibition onto MCs, these cells may be required to fill multiple functional roles, providing regulation of both temporal precision, which is canonically driven by feedforward PV input, and gain modulation, a role traditionally associated with CCK interneurons. Our findings indicate that VGluT3+ interneurons can be engaged in both feedforward and feedback circuits, which, together with their distinct synaptic properties, may enable VGluT3+ interneurons to both influence temporal precision in MCs and integrate information over longer time scales to modulate MC output. However, more work is needed to precisely discern how VGluT3+ interneurons in DG impact excitability and input-output transformations of MCs in these pathways.

Interestingly, we found that postsynaptic target selectivity was a property not only of VGluT3+ interneurons but also PV, SOM, VIP and VGluT3-/CCK+ interneurons in DG. This differential organization of inhibitory inputs onto MCs and GCs is a circuit feature that can facilitate the disparate activity patterns observed in MCs and GCs in vivo as MCs receive minimal input from the inhibitory sources they recruit to dampen GC activity ^13–15,18^. In addition, the primary source of inhibition onto MCs is CB1+ and exhibits cannabinoid-mediated short-term plasticity indicating that MC activity can lead to disinhibition through retrograde endocannabinoid signaling, providing another mechanism to maintain high MC activity levels. The ability to independently regulate MC excitability supports theoretical and computational work positing that MCs contribute to pattern separation in the DG through regulation of GC sparsity^13,45,46^. Indeed, ablation of MCs leads to transient GC hyperexcitability and deficits in contextual discrimination^21^. Further corroborating this circuit dynamic, we found that VGluT3+ interneuron-mediated alterations in MC activity led to opposing changes in GC activity indicating that at a population level MCs drive inhibition of GCs. We also found that manipulating VGluT3+ interneurons altered the spatial tuning fraction of MCs and remapping of MCs suggesting selective inhibitory circuits contribute to DG spatial representations and context discrimination. Taken together, our findings provide further support for the combined role of MCs and GCs in pattern separation, with high MC activity serving to constrain GC representations, and offer a circuit level mechanism that enables this dynamic.

Our findings on the evolutionary conservation of inhibitory connectivity in DG adds to a growing body of work examining the translational relevance of rodent findings to the human brain (see ^47–55^). In addition, these findings may provide insight into circuit maladaptations found in temporal lobe epilepsy (TLE), which has previously been linked to MC disruption^22,46,56–58^. We recently found that hilar excitatory cells, including MCs as well as CA3 and CA4 pyramidal cells (PCs), in tissue resected from pharmacoresistant epilepsy patients had excitation/inhibition imbalances and dramatic reductions in inhibitory input when compared to rodents^47^. In these experiments, we found that stimulation intensities that were sufficient to drive excitatory mossy fiber input onto hilar excitatory cells were unable to recruit concomitant disynaptic inhibition. However, in the current study, extracellular stimulation at the hilar/GCL border was still able to drive monosynaptic inhibition onto MCs suggesting E/I imbalances at hilar excitatory cells may not reflect complete loss of inhibitory input but rather deficits in circuit recruitment of inhibition, particularly that mediated by VGluT3+ interneurons, or alterations in interneuron excitability. Alternatively, as VGluT3+ interneurons co-release glutamate and compromised glutamate decarboxylase (GAD) function can lead VGluT3+ interneurons to exert a net excitatory influence on the hippocampal network, it’s also possible that circuit conditions in TLE may shift synaptic output in favor of glutamate, reducing overall inhibitory tone^25^. Future studies exploring the synaptic properties and circuit integration of VGluT3+ interneurons in epileptic conditions will help delineate how these alterations in VgluT3+ interneuron function may contribute to circuit hyperexcitability.

## Acknowledgements

Supported by an NICHD Intramural Research Program (IRP) grant awarded to C.J.M., NINDS IRP grant to K.Z., NIMH IRP grant to B.B.A., and NIH Center on Compulsive Behaviors (CCB) fellowship to G.A.V. and A.P.C. Work was carried out in collaboration with the NIH Comparative Brain Physiology Consortium (CBPC).

## Contributions

G.A.V was responsible for conceptualization, experimental design, IHC, electrophysiology, multiphoton imaging, data analysis and writing. K.A.P. and C.J.M. contributed to project conceptualization, experimental design, supervision and editing of manuscript. H.R. and X.Y. contributed to IHC and RNAscope experiments. B.E.H., A.P., A.M., M.E. and B.A. conducted nonhuman primate injections. K.Z. performed human patient resections. E.F., Y.W., M.D., B.L.G., J.D., and G.F. contributed design and production of novel enhancer viruses.

## Methods

### Experimental Models and Subjects

#### Mice

Experiments were performed in male and female VgluT3^Cre^, PV^Cre^, SOM^Cre^, VIP^Cre^, 5HT3-GFP and SNCG^Flpe^ transgenic mice. Animals ranged in age from postnatal day 45 (p45)-p90 for electrophysiology and immunohistochemistry experiments and from p60-p150 for in vivo 2-photon imaging experiments. For targeted electrophysiological recordings Cre mouse lines were crossed with tdTomato^fl/fl^ (Ai14) mice. Channelrhodopsin experiments were performed by crossing Cre lines with ChR2^fl/fl^ (Ai32) mice. All mouse lines were maintained on a C57/B6 background. Mice were housed with a 12hr light/dark cycle and fed standard rodent chow. All rodent experiments were conducted in accordance with an Animal Study Protocol approved by the ACUC at the National Institute of Child Health and Human Development.

#### Nonhuman Primates

Rhesus macaque tissue was obtained through the NIH Comparative Brain Physiology Consortium (CBPC) from 7 macaques (2 female, 5 male) aged 9-15 years. All experiments were conducted in accordance with the ILAR Guide for Care and Use of Laboratory Animals and were conducted under an Animal Study Protocol approved by the ACUC at the National Institute of Mental Health. All procedures adhered to the applicable Federal and local laws, regulations and standards including the Animals Welfare Act and Public Health Service policy (PHS2002). To acquire tissue for acute slice preparation, the animals were sedated with a combination of ketamine (5-15mg/kg) and midazolam (0.05-0.3mg/kg) and anesthetized with inhaled isoflurane. Prior to brain removal and blocking, animals were transcardially perfused with ice-cold sucrose-substituted artificial cerebrospinal fluid (ssaCSF) containing (in mM): 90 sucrose, 80 NaCl, 24 NaHCO_3_, 10 glucose, 3.5 KCl, 1.25 NaH_2_PO_4_, 4.5 MgCl_2_ and 0.5 CaCl_2_ saturated with 95% O_2_ and 5% CO_2_.

For viral expression in NHPs, AAVs containing ChR2-mCherry fusion protein under control of the Dlx, s5e2 or BiSSTE10 promoter (AAV.PhP.Eb-mDlx-ChR2-mCherry, AAV.PhP.Eb-s5e2-ChR2-mCherry or AAV.PhP.Eb-BiSSTE10-ChR2-mCherry) were injected in rhesus macaques. In each hemisphere of each subject, 20-25μL of virus was injected at each of two anterior-posterior locations spaced approximately 2mm apart, and caudal to the uncus. Stereotaxic coordinates for injections were derived from MRI and virus was delivered using a needle guide for enhanced accuracy^59^. Surgeries were performed under aseptic conditions in a fully equipped operating suite.

#### Human Tissue

For electrophysiological recordings from human tissue, tissue was surgically resected from XX anonymized/deidentified participants with pharmacoresistant epilepsy. The participants underwent an initial surgical procedure during which recording electrodes were implanted subdurally on the cortical surface and within the brain parenchyma to monitor epileptiform activity. The location of the intracranial electrodes was selected by the clinical team to localize the epileptogenic zone and recordings during a monitoring period were used to identify the specific hippocampal region exhibiting ictal or inter-ictal activity. During a second surgery the brain areas of seizure onset were surgically resected. The Institutional Review Board (IRB) at the National Institute of Neurological Disease and Stroke approved the research protocol and informed consent for the experimental use of surgically resected tissue was obtained from each participant and their guardian.

## Method Details

### Immunohistochemistry

Mice were transcardially perfused with 4% PFA and dissected brains were placed in 4% PFA for 24 hours. Brains were then washed with 1X PBS and cryoprotected with 30% sucrose in PBS. 45 μm sections were cut using a freezing microtome (Mircom, Waltham, MA) and sections were placed in 1x PBS. Free floating sections were then blocked and permeabilized with Blocking solution (1x PBS, 10% normal goat serum, 0.5% Triton X-100) at room temperature for at least 2 hours. Following blocking, primary antibodies were diluted in carrier solution (1x PBS. 1% normal goat serum, 0.1% Triton X-100) and sections were incubation with primary antibodies at 4° C for 48 hours. Brain slices were then washed with 1x PBS for 15 minutes 4 times before being incubated with secondary antibodies diluted in carrier solution at room temperature for 1 hour. Following 4 washes with 1x PBS sections were then mounted on gelatin coated slides, air dried and cover slipped with Prolong Diamond Antifade mountant with DAPI (ThermoFisher Scientific Cat. No. 36962). Slides were cured at room temperature before imaging.

### Morphological reconstruction

Brain sections containing biocytin-filled cells were drop fixed overnight in 4% PFA at 4° C immediately after electrophysiological recording. Sections were then washed in 1x PBS, permeabilized with 0.3% Triton X-100 and incubated overnight Alexa-488, Alexa-555 or Alexa-647 conjugated streptavidin overnight (ThermoFisher Scientific Cat. No. S11223, S21381, S21374). If additional staining was desired, sections were blocked and permeabilized with blocking solution for two hours at room temperature. Sections were then incubated with primary antibodies diluted in carrier solution for 3 hours at room temperature. Following 4 washes with 1x PBS for 15 minutes each, sections were incubated with secondary antibodies diluted in carrier solution at room temperature for 2 hours. Sections were then incubated with 1μg/mL DAPI (Millipore Sigma Cat. No. D9542) before undergoing 4 washes with 1x PBS and cryopreserved in 30% sucrose. Brain slices were then resectioned into 70 μm sections using a freezing microtome (Microm, Waltham, MA) and mounted on gelatin coated slides. Sections were cover slipped with Mowiol mounting media and cured at room temperature before imaging.

### In situ hybridization (ISH) RNAScope

Tissue blocks dissected from macaques were submerged in 2-methylbutane that was pre-chilled in a dry ice/ethanol bath for 1 minute. Tissue was removed, wrapped in foil and stored at -80° C. 10 μm sections were cut using a Leica Cryostat and transferred to gelatin coated slides. Slides were stored at 80° C until processing.

Probes were designed and manufacture by ACD Bio. ISH was carried out following the RNAScope Multiplex Fluorescent v2 Assay developed by ACD Bio. Briefly, slide mounted sections were fixed with 4% PFA and dehydrated with increasing concentrations of ethanol (50%, 70% and 100%). The slides were then treated with H_2_O_2_ and Protease IV before incubation with probes for 2 hours at 40° C. Probes were detected using the RNAScope Multiplex Detection v2 kit. Sections were stained with DAPI and cover slipped with Prolong Gold Antifade mountant (ThermoFisher Scientific Cat. No. 10144).

### IHC and ISH imaging and analysis

Fluorescent IHC images were acquired on a Zeiss LSM 900 confocal microscope. Images of axonal boutons were acquired using airyscan at 63x and images for morphological reconstructions were acquired at 20x. For each experiment, microscope settings – laser power, gain, offset, averaging, pixel density and optical section thickness – were kept the same. Images were acquired as a tiled z-stack with 10% overlap between tiles and 50% overlap for each optical section in z. RNAScope images were acquired using with an SLIDEVIEW VS200 slide scanner (Olympus) with an ORCA-Fusion CMOS camera (Hamamatsu).

Morphological reconstructions were performed on z-stacks using the simple neurite tracer plugin for ImageJ, a semiautomated segmentation function. Visually identified dendrites and axons were separately traced. Profile plots of axonal bouton intensity were generated using the plot profile function in ImageJ.

### Electrophysiology

#### Acute Slice Preparation

Adult mice were anesthetized using isoflurane and rapidly decapitated. Brains were removed, blocked and sectioned in ice-cold choline cutting solution containing (in mM) 110 choline chloride, 25 NaHCO_3_, 25 glucose, 11.6 Na-Ascorbate, 3.1 Na-Pyruvate, 2.5 KCl, 1.25 NaH_2_CO_4_, 7 MgCl_2_ and 0.5 CaCl_2_ saturated with 95% O_2_ and 5% CO_2_. 300 μm horizontal sections were cut using a VT-1200s vibratome (Leica Microsystems) and transferred to a submerged incubation containing warmed (32-34° C) choline cutting solution. After 15 minutes slices were transferred a submerged incubation chamber containing room temperature high-Mg^2+^ aCSF containing (in mM) 130 NaCl, 24 NaHCO_3_, 10 glucose, 3.5 KCl, 1.25 NaH_2_PO_4_, 5 MgCl_2_ and 1 CaCl_2_ with osmolarity of 300-310 and saturated with 95% O_2_ and 5% CO_2_ for at least 45 minutes before recording. Sections were kept at room temperature for the remainder of the day.

For macaque and human tissue, tissue blocks were trimmed and sectioned with a VT-1200S vibratome in ice-cold ssaCSF. After sectioning, slices were transferred to a submerged incubation chamber containing warmed (32-34° C) ssaCSF for 30 min and then maintained at room temperature for up to 48 hours with solution changes every 24 hours.

#### Electrophysiological Recording

After incubation slices were transferred to a submerged recording chamber. Slices were constantly perfused with oxygenated aCSF containing (in mM) 130 NaCl, 24 NaHCO_3_, 10 glucose, 3.5 KCl, 1.25 NaH_2_PO_4_, 1.5 MgCl_2_ and 2.5 CaCl_2_ with osmolarity of 300-310 at a temperature of 32-34° C and a flow rate of 2-3 mL/min. As required aCSF was supplemented with 10 μM bicuculline methobromide (Tocris Cat No. 0109) or 5 μM WIN55,212-2 (Tocris Cat No. XXXX). Individual cells were visualized with a 40x objective with IR-DIC video microscopy under an upright microscope (Zeiss Axioscope) and fluorescent labelled cells were identified using widefield LED illumination (CoolLED) through the objective. Whole cell patch clamp recordings were performed on slices with electrodes (3-5 MΩ) pulled from borsillicate glass (World Precision Instruments, Sarasota, FL, USA) filled intracellular recording solution (ICS). Field recording electrodes were filled with aCSF. Two different ICS’ were used. For intrinsic electrophysiological properties and presynaptic cells in paired recordings ICS contained (in mM) 130 K-gluconate, 10 HEPES, 5 KCl, 3 MgCl_2_, 2 Na_2_ATP, 0.3 NaGTP, 0.6 EGTA and 0.3-0.3% biocytin. For optogenetic, evoked IPSC and postsynaptic cells in paired recordings ICS contained (in mM) 130 CsCl_2_, 8.5 NaCl, 5 HEPES, 4 MgCl_2_, 4 Na_2_ATP, 0.3 NaGTP, 1 QX-314 and 0.2-0.3% biocytin (E_Cl_ = 0mV). ICS was adjusted to a pH of 7.2-7.3 and total osmolarity of 290mM. Voltage and Current-clamp recordings were recording using a Multiclamp 700b amplifier (Molecular Devices, Sunnyvale, CA, USA) and signals were digitized with a Digidata 1440 board (Molecular Devices, Sunnyvale, CA, USA) at 10-20 kHz. Signals were filtered at 4-10kHz (Bessel) and acquired using pClamp10.7 software (Molecular Devices, Sunnyvale, CA, USA). For all recordings the fast capacitance transient was removed using digital leak subtraction and uncompensated series resistance (5-15 MΩ) was monitored throughout recordings. Recordings were discarded if series resistance changed >10%. Membrane potentials were not corrected for liquid junction potentials.

Intrinsic electrophysiological properties were determined as previously described^60^. Briefly, resting membrane potential was measured through cell-attached voltage clamp recordings. A 100ms voltage ramp was applied from 100mV to -200mV and the K^+^ reversal potential was determined after subtracting the K^+^ leak current^61^. Action potential threshold, after-hyperpolarization amplitude, action potential amplitude and action potential kinetics were measured in current clamp by applying a 1s voltage step. Measurements were taken from the first spike at rheobase. Threshold was determined as the voltage at which dV/dT exceeds 10mV/ms and max upstroke and downstroke were determined as the maximum and minimum of the first derivative of the spike waveform, respectively. Sag ratio, an indirect measure of I_h_, was calculated as the steady state voltage divided by the peak voltage in response to a 1s hyperpolarizing current step that drove the cell to -100mV. Maximum firing frequency is reported as the maximum sustained firing frequency recorded in response to 1s depolarizing current steps. Accommodation ratio was measured at maximum firing frequency and calculated as the first interspike interval divided by the average of the last two interspike intervals. Input resistance was measured by recording the voltage in response to 20 current steps starting from -50 mV in 5 mV increments. Membrane time constant (tau) was measure by taking the average of 20 sweeps of a 400ms current step to -20 mV. Tau was measured by fitting a single exponential to the voltage waveform at the start of the current step.

For paired recordings a minimum of 10 trials were analyzed to acquire unitary IPSC properties and measure synchronicity. For each recording a train of 25 presynaptic action potentials were delivered at 50 Hz by applying 1-2 nA current steps of 1-2 ms duration to the presynaptic cell while postsynaptic cells were held -70 mV. uIPSC amplitude was measured as the maximum postsynaptic response, including failures, during the first AP in the train. Paired pulse ratio was calculated by dividing the amplitude of the uIPSC during the second AP by the first uIPSC in the 50 Hz train. Latency was measured as the time from the peak of the presynaptic spike to the start of the postsynaptic uIPSC. Asynchronous release was measured by deconvolution analysis as previously described ^25,31^. Briefly, an artificial miniature IPSC (mIPSC) was created using the rise time and decay kinetics (measured by a single exponential) of a single unitary response and scaled 20 pA. Postsynaptic responses during a 50 Hz train were smoothed by 20 repetitions of Gaussian smoothing. Fast fourier transforms (FFT) were then performed on the mIPSC waveform and postsynaptic response and the mIPSC FFT was divided point-by-point into the FFT of the postsynaptic waveform. The result of this division was then converted back into the time domain by inverse FFT, creating a release rate histogram (RRH). Synchronicity ratio was calculated by taking the area under the curve (AUC) of the RRH with synchronous release being defined as the AUC during the 5 ms window immediately following the onset of the presynaptic current step and asynchronous as the AUC during 15 ms window preceding the onset of the next current step. The synchronicity ratio represents synchronous release/asynchronous release for each presynaptic current step.

For stimulation experiments, electrical stimulation was applied by a constant current isolation unit (A360, World Precision Instruments) connected to either a glass electrode filled with aCSF or concentric bipolar electrode. Monosynaptic inhibition was evoked by placing a glass stimulating electrode in the hilus just inside the GCL. For feedforward pathway experiments, excitation was evoked by placing a bipolar electrode in the middle molecular layer. In both experiments low intensity (15-120 μA) stimulation was applied and adjusted until electrical stimulation reliably evoked a postsynaptic event (e.g. IPSC or field EPSP) without failures (∼50% of maximum response). DSI was induced by depolarizing the postsynaptic cell from -70 mV to 0 mV for 2s and measured as the change in amplitude of the eIPSC recorded immediately preceding the depolarization to one just after the depolarization.

For optogenetic experiments, light was delivered through a 40x objective. Light wavelength and intensity were regulated by a CoolLED PE-4000(Andover, United Kingdom) with 480 nm light used to activate ChR2. Cells recorded during simultaneous patch clamp experiments were <100 μm apart and the objective was centered over the cell pair for light stimulation.

### Stereotaxic injections and cranial window implant

#### Viral constructs

AAV2-retro-hSyn-mCherry (Addgene, Cat. No. 114472-AAVrg) and AAV9-CAG-flex-EGFP (Addgene, Cat. No. 51502-AAV9) viruses were used to label MCs and VGluT3+ interneurons in VGluT3^Cre^ mice. AAV2-retro-hSyn-ChR2-EYFP (Addgene, Cat. No. 26973-AAVrg) was used to express ChR2 in MCs. The novel enhancer viruses AAV.PHP.eB-mDlx-ChR2-mCherry, AAV.PHP.eB-s5e2-ChR2-mCherry (Addgene, Cat. No. 135634-PHPeB) and AAV.PHP.eB-BiSSTE10-ChR2-mCherry (Addgene, Cat. No. 213815-PHPeB) were used in NHPs to selectively drive ChR2 expression in Dlx+ interneurons, PV+ interneurons and SOM+ interneurons, respectively^34–36^. AAV9-CaMKII-GCaMP6f (Addgene Cat. No., 100834-AAV9) and AAV5-CaMKII-GCaMP8f (Addgene Cat. No., 176750-AAV5) viruses were used to express GCaMP in GCs and MCs. AAV9-hSyn-DIO-hM3D(Gq)-mCherry (Addgene Cat. No., 44361-AAV9)and AAV9-hSyn-DIO-hM4Di-mCherry (Addgene Cat. No., 44362-AAV9) were used in VGluT3^Cre^ mice to drive expression of DREADDs in VGluT3+ interneurons

#### Rodent Injections

Mice were induced with 5% isoflurane, mounted in the stereotax and maintained with 1.5-2.5% isoflurane for the duration of the surgery. Prior to the start of the surgery mice were given buprenorphrine (0.1 mg/kg) for post operative analgesia. Viruses were injected into left dorsal DG (AP: -2.1mm ML: -1.45mm DV: -1.95, -2.05mm) for retrograde MC labeling, right dorsal DG (AP: -2.1, -2.3mm, ML: 1.45mm, DV: -1.95, -2.05mm) for GCaMP/DREADD experiments and bilaterally in ventral DG (AP: -3.65mm, ML: ± 2.6mm, DV: -2.75, -3.10, -3.5mm) for enhancer virus experiments. 80 nL of virus was injected at each location at a rate of 100 nL/min using a Nanoinjector (Neurostar, Germany). Following surgery mice were given ketoprofen and topical triple antibiotic ointment with lidocaine for 3 days.

#### Cranial Window

One week after GCaMP/DREADD injections mice were implanted with a cranial window and stainless steel headbar for head fixation during imaging. Imaging cannulas for DG access were created by adhering a 3 mm circular coverslip (World Precision Instruments) to a cylindrical stainless steel cannular (diameter: 3 mm, height: 1.5 mm) using optical adhesive (Norland). The imaging cannula was placed ∼100 μm above the hippocampal fissure providing optical access to the dorsal blade of the GCL and the hilus. A circular headbar was implanted on the skull surrounding the cannula. Mice were anesthetized with isoflurane (induction: 5%, maintenance: 1.5-2.5%) and given buprenorphrine. The scalp was them removed a 3 mm craniotomy over right DG was performed with a robotic drill (Neurostar, Germany). The dura was removed, and underlying cortex was slowly aspirated using a blunted 25 g needle. After reaching the fibers of the external capsule, a 27 g blunt needle was used to slowly aspirate CA1 until the loose fibers and vasculature of stratum lacunosum moleculare was visible. Bleeding was controlled with hemostatic collagen sponges (Surgifoam, Johnson and Johnson) soaked in ice-cold aCSF and constant irrigation with ice-cold aCSF. The imaging cannula was then inserted and a headbar was attached to the skull with Metabond dental cement (Parkell). Mice were then allowed to recover in their home cage placed over heating pad and were given ketoprofen and topical triple antibiotic ointment for three days postoperatively.

### Virtual environment and behavioral training

Mice were allowed to recover for 1 week after window implantation before starting behavioral training. Mice were first habituated to handling and head-fixation while on top of a floating Styrofoam ball with motion restricted to forward-backward running, progressively increasing the duration of head fixation from 5 minutes to 20 minutes over 4-5 days. After habituation to head-fixation, mice were introduced to a linear virtual reality environment (Familiar environment) with optical sensors on the styrofoam ball translating the mouse’s movement into the virtual space. The environment was projected onto a dome (Phenosys, Germany) that surrounded ∼90% of the mouse’s visual field and was created with Phenosys’ proprietary software, consisting of textured walls, floors and 3D rendered objects placed along the sides of the track to serve as visual contextual cues. Once mice reached the end of the linear environment they were teleported back to the beginning of the track after a brief (1s) pause. During initial training runs sucrose rewards were dispensed through a lickport at random locations throughout the maze. Once the mice showed consistent running (∼1 lap every 2 minutes), rewards were then restricted to a specific location along the maze. During training sessions mice were also habituated to the optical instrumentation (i.e. overhead objective, laser and shutter noises) prior to sessions used for image acquisition. Imaging sessions for single environment experiments had a duration of 15 mins in the familiar environment. For contextual discrimination sessions, mice were randomly teleported at the end of each run to the familiar environment or a novel environment that contained distinct textures on the walls, different 3D visual cues along the track and a different reward location. Imaging sessions for contextual discrimination experiments were 20 minutes in duration.

### Chemogenetic manipulation of VGluT3+ interneurons

During behavioral training mice were habituated to intraperitoneal (i.p.) injections, receiving i.p. injections of saline prior to randomly selected training sessions. For both hM3Dq and hM4Di experiments, mice received i.p. injections of DREADD agonist 21 (C21) (3 mg/kg; Tocris Cat No. 6422) 30 minutes before the start of imaging sessions. Mice were randomly assigned to two groups, with one group undergoing baseline imaging 24 hours before C21 imaging sessions while the other group’s baseline imaging session took place 24 hours after C21 imaging sessions. This counterbalanced design was implemented to mitigate any sequence effects when comparing baseline to C21 imaging sessions.

### Two-photon Imaging

Images were acquired with a rotating 8 kHz galvo-resonant laser scanning 2-photon microscope (Bergamo, Thor Labs) at a frame rate of 30Hz using bidirectional scanning. Images were acquired through a 10x objective (Thor Labs, 0.5 N.A., 7 mm WD) that was rotated so that the objective was parallel with the imaging window to optimize light transmission. GCaMP6f/8f was excited at 920 nm with a femtosecond tunable laser (MaiTai DeepSee, Spectra-Physics). Laser power was 30-80mW as measured under the objective with power level varying depending on depth of the imaging plane and window clarity. Emitted signals were captured with a photomultiplier tube (GaAsP PMT, Thor Labs) at 512×512 pixels with field of view covering 619 μm x 619 μm for MC imaging and 400 μm x 400 μm for GC imaging. To block ambient light from the virtual reality system, a custom designed, flexible 3D printed cone was placed around the objective, forming a seal with the circular headbar.

### 2-photon data processing and transient detection

Raw movies were first motion-corrected using suite2p^62^. Putative cells were then segmented using suite2p automated segmentation. ROIs identified through automated segmentation were then manually curated using the suite2p GUI. MCs and GCs were identified by their somatic location in the DG and morphology with MCs having large, multipolar cell bodies located in the hilus while GCs were characterized by small somas densely packed in the GCL.

Fluorescence signals extracted from ROIs were baseline corrected and converted to relative fluorescence change (ΔF/F) as described in Jia et al.^63^ with a uniform smoothing window of t_1_ = 3 s and baseline window t_2_ = 60 s. Putative calcium transients (>2 standard deviations from total mean fluorescence) were removed prior to baseline correction and ΔF/F conversion. This process was repeated twice to remove potential transient contamination from baseline determination.

Significant calcium transients were identified by a threshold approach based on amplitude and duration as previously described^13,64^. Potential transients were initially identified as events that exceed 3 s.d. with event onset being defined as the frame when the signal exceeds this value and event offset defined as the frame when the signal subsequently falls below 0.5 s.d. Significant transients were then identified as those events with a false positive rate <5% based on duration with the false positive rate determined by the ratio of negative to positive going transients with a given duration. For analysis of transient frequency only cells that had at least one significant transient during the imaging session were included.

### Spatial tuning

To examine MC spatial tuning, we first created a spatial tuning curve for each cell by breaking the 3m long virtual environment into 100 3 cm-wide bins. For each spatial bin, we then calculated the average ΔF/F, including only frames where the mouse was running (>1 cm/s) and smoothed the resulting curve with a Gaussian filter (σ = 3) to create the spatial tuning curve. Only mice that ran more than 5 laps in each environment were included in spatial tuning analysis.

To determine if a cell was spatially tuned, we created a null tuning distribution by circularly rotating position relative to ΔF/F and recreating the spatial tuning curve, repeating this procedure 1,000 times. A cell was then determined to be spatially tuned if it had at least 5 consecutive spatial bins (15 cm) that exceeded the 95^th^ percentile of the shuffled distribution and had a transient within the field on at least 25% of laps. Place field width was calculated as the number of consecutive spatial bins that exceeded that half maximum of the spatial tuning curve.

### Activity and velocity cross-correlation

To calculate the correlation between each MCs activity and the mouse’s running velocity, we shifted the fluorescence trace relative to the smoothed velocity trace one frame at a time from -5 s to +5 s and calculated the Pearson’s correlation coefficient at each temporal shift. The maximum absolute value of all the cross correlations within this temporal range was taken as the correlation coefficient for each cell and the shift where this value occurred was taken as the lag with negative lag values indicating activity trails velocity while positive values indicate activity leads velocity.

### Remapping analysis

For MCs that were tuned to at least one of the two contexts (familiar or novel), we calculated the correlation of tuning within the familiar context and between the familiar/novel contexts. To calculate the within context correlations, we split the laps run in the familiar context in half and created a spatial tuning curve for each half. The spatial tuning correlation was then calculated as the Pearson’s correlation coefficient between the two spatial tuning curves for each tuned cell. For between context correlations, we created spatial tuning curves for each context and took the Pearson’s correlation coefficient between familiar and novel context tuning curves. Change in stability was calculated as the difference between the within context correlation coefficient and the correlation coefficient across contexts (r_9_ – r_fn_).

### Statistics

Statistical tests were performed in either GraphPad Prism (GraphPad Software, San Diego, CA, USA) or with custom Python code. All datasets that underwent statistical testing were first tested for normality using a Shapiro-Wilke test. Following determination of normality, the appropriate parametric or non-parametric statistical tests were used to assess statistical significance. All tests were two-sided. Throughout group data are presented as mean±s.e.m. with symbols representing individual values unless otherwise specified in figure legends. For box plots, horizontal lines represent the median with boxes representing 25^th^-75^th^ percentile and whiskers representing 1.5 times the interquartile range.

---

**Supplemental Figure 1.**
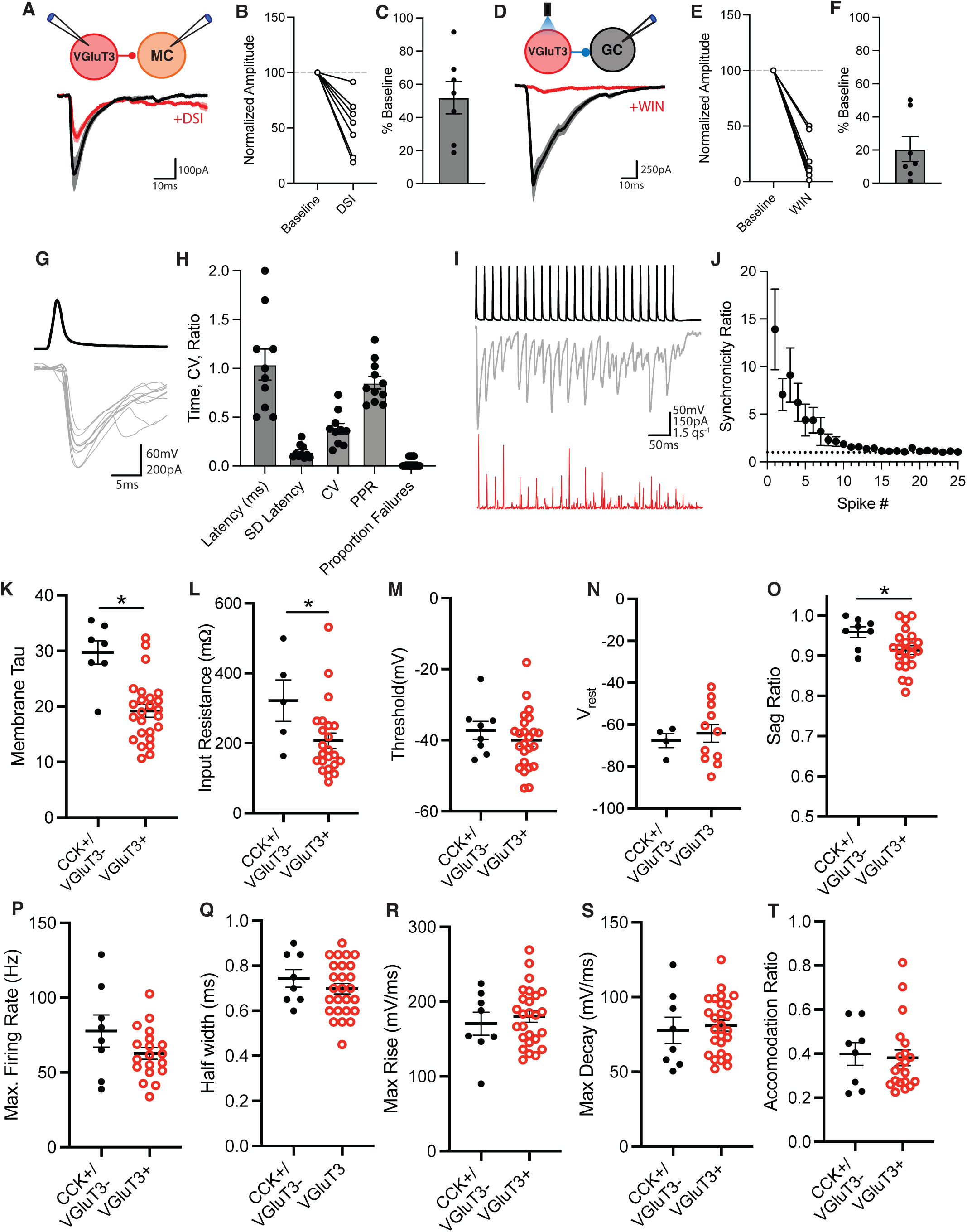
Synaptic properties of DG VGluT3+ interneurons. **A)** Schematic of recording setup for paired VGluT3+ interneuron-MC recordings (top) and representative traces of uIPSC at baseline and after DSI (bottom). **B)** Graph showing normalized amplitude (as % of baseline) during baseline and after DSI. **C)** Group data showing average amplitude as a percentage of baseline after DSI (n=7 pairs, 3 mice; mean± s.e.m.). **D)** Schematic of recording setup for examining VGluT3+ interneuron leIPSCs simultaneously in MCs and GCs (top) and representative traces of leIPSCs in MCs during baseline and after bath application of WIN55,212-2 (bottom). **E)** Plot showing normalized amplitude (as % of baseline) during baseline and after WIN application. **F)** Group data displaying average leIPSC amplitude in MCs after WIN expressed as percentage of baseline (n=7 cells, 2 mice; mean± s.e.m.). **G)** Representative traces of VGluT3+ mediated uIPSC recorded in an MC. **H)** Group data on VGluT3+ interneuron synaptic latency, jitter (SD of latency), coefficient of variation (CV), paired-pulse ratio (PPR) and proportion of failures recorded in postsynaptic MC (n=10 pairs, 5 mice; mean± s.e.m.). **I)** Representative traces of presynaptic AP train in VGluT3+ interneuron (top), postsynaptic uIPSCs in MC during the train (middle) and release rate histogram (bottom). **J)** Plot show synchronicitiy ratio of VGluT3+ interneuron mediated uIPSCs across a 25-pulse train at 50 Hz. Dotted line indicates SR of 1 (n=10 cells, 5 mice). **K-T)** Plots showing passive and active electrophysiological properties for CCK+/VGluT3- and VGluT3+ interneurons in DG (*<0.05, t-test, mean± s.e.m.).

**Supplemental Figure 2.**
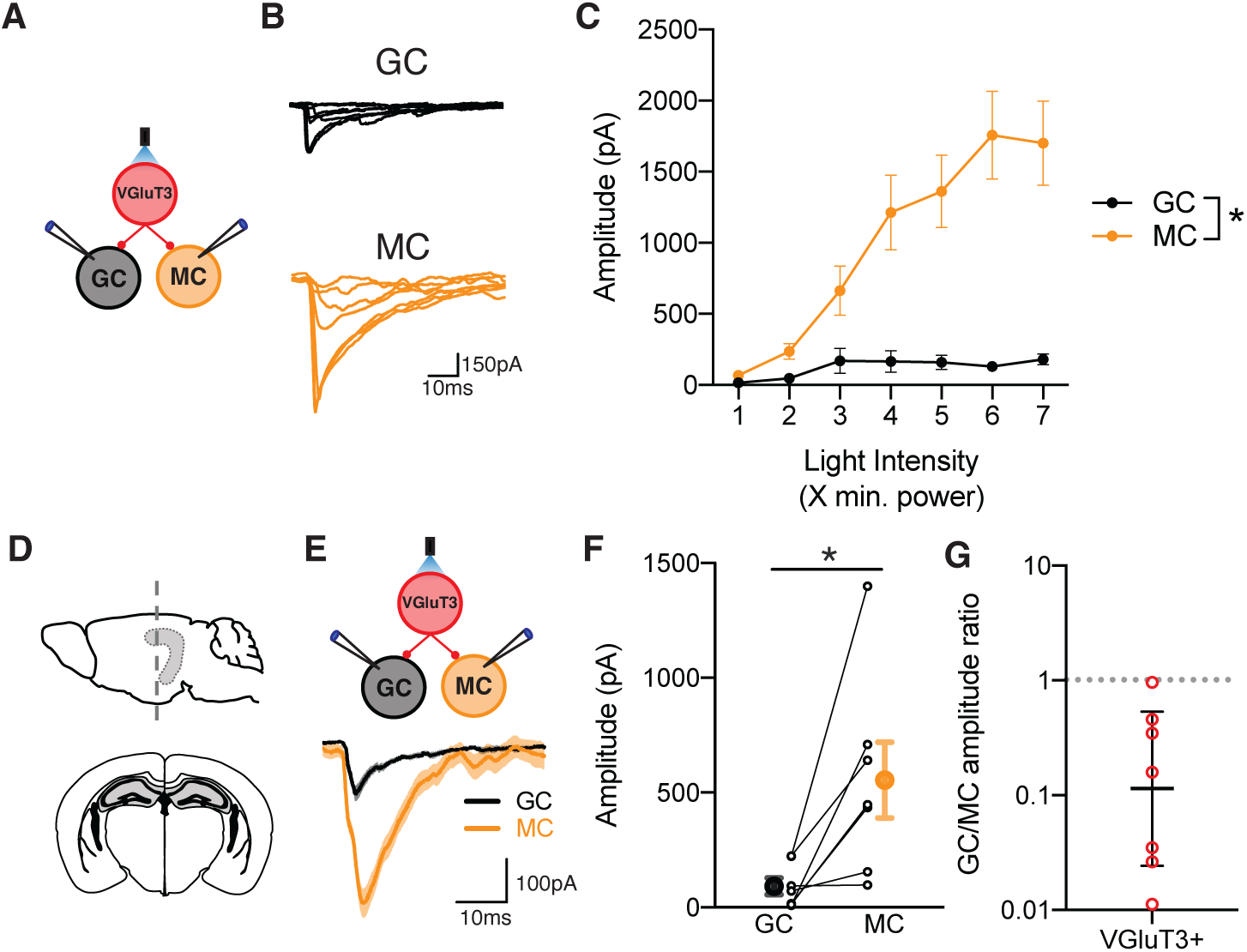
Optogenetic input-output relationship for VGluT3+ interneurons and VGluT3+ target selectivity in dorsal DG. **A)** Diagram of recording setup for recording VGluT3+ interneuron leIPSCs simultaneously in MCs and GCs. **B)** Representative traces of leIPSCs recording GCs (top) and MCs (bottom) across different light intensities. **C)** Plot showing average leIPSC in GCs and MCs at different light intensities. Y axis represents fold change in intensity from minimum photostimulation intensity (GC, n=12 cells, 3 mice; MC, n=14 cells, 3 mice; *p<0.05 across all points – multiple t-tests; mean± s.e.m.). **D)** Schematic of coronal sectioning plane to record from dorsal DG. **E)** Diagram of recording setup for optogenetic stimulation of VGluT3+ interneurons in dDG (top) and representative traces of leIPSCs recorded in GCs and MCs (bottom). **F)** Group data showing leIPSC amplitude in GCs and MCs, mean± s.e.m.. **G)** Plot showing leIPSC amplitude ratio (GC/MC) for each GC and MC pair – geometric mean ± s.e.m. (n=7 pairs, 2 mice; *p<0.05 paired t-test).

**Supplemental Figure 3.**
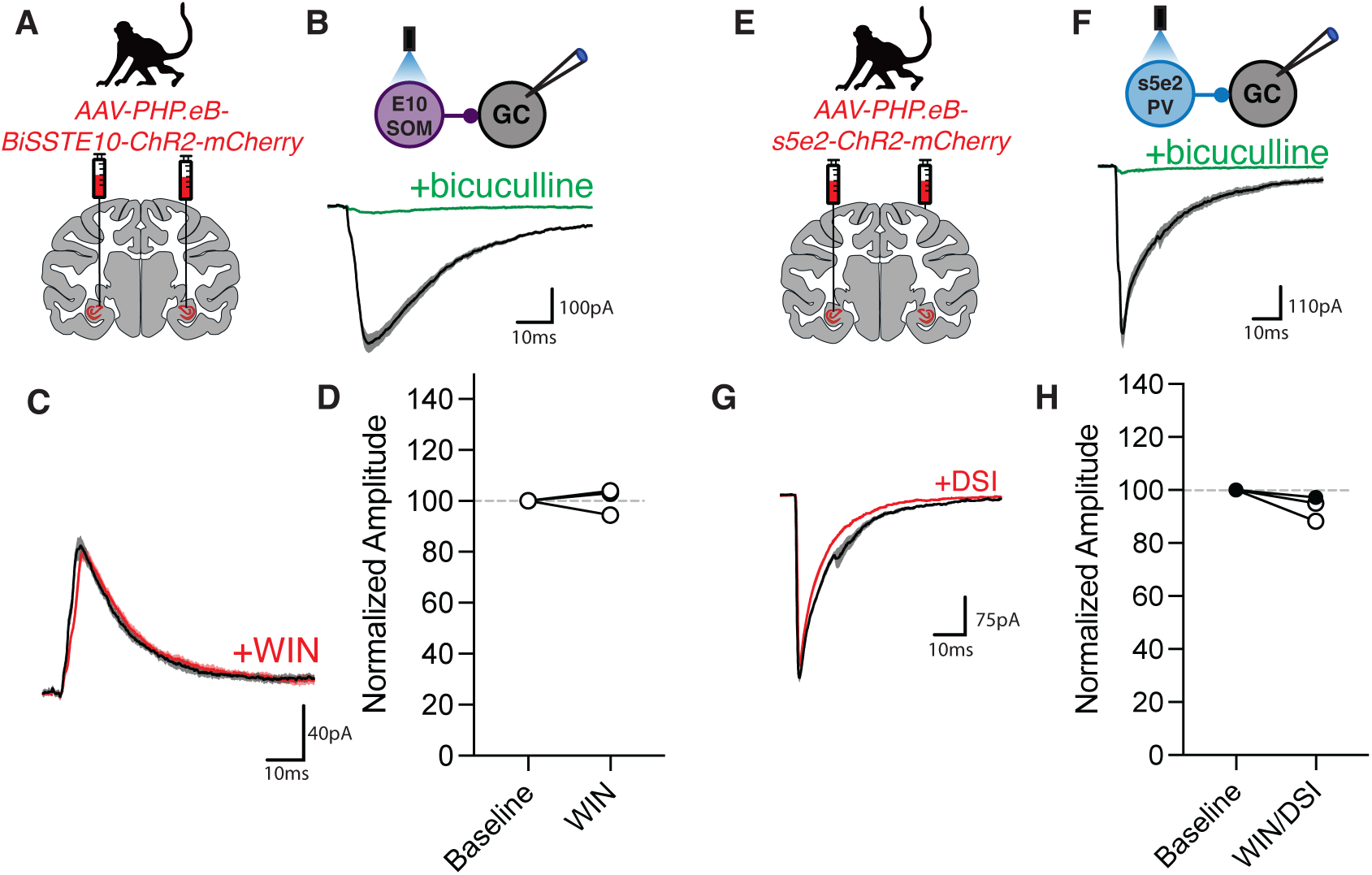
Pharmacological characterization of BiSSTE10 and s5e2 leIPSCs in non-human primates. **A)** Schematic of bilateral virus injection of BiSSTE10-ChR2-mCherry in macaque hippocampus. **B)** Diagram of recording setup (top) and representative traces showing that BiSSTE10 leIPSCs are blocked by bath application of the GABA-A receptor antagonist bicuculline (bottom). **C)** Representative traces of leIPSCs mediated by BiSSTE10+ cells recorded in GCs during baseline and after DSI induction. Note BiSSTE10 leIPSCs are not cannabinoid sensitive. **D)** Plot showing normalized (as % of baseline) BiSSTE10 leIPSC amplitude during baseline and after DSI induction. Note these recordings were with V_hold_=-30mV and E_Cl_=-90mV. **E)** Same as in A for s5e2-ChR2-mCherry virus injection. **F)** Recording diagram (top) representative traces of s5e2 leIPSCs recorded in GCs during baseline and after bath application of bicuculline (bottom). **G)** Representative traces of s5e2 leIPSCs during baseline and after induction of DSI. As with BiSSTE10 note the lack of cannabinoid sensitivity. **H)** Plot showing normalized (as % of baseline) s5e2 leIPSC amplitude during baseline and after DSI induction or bath application of WIN55,212-2. Filled circles indicate WIN and open indicate DSI.

**Supplemental figure 4.**
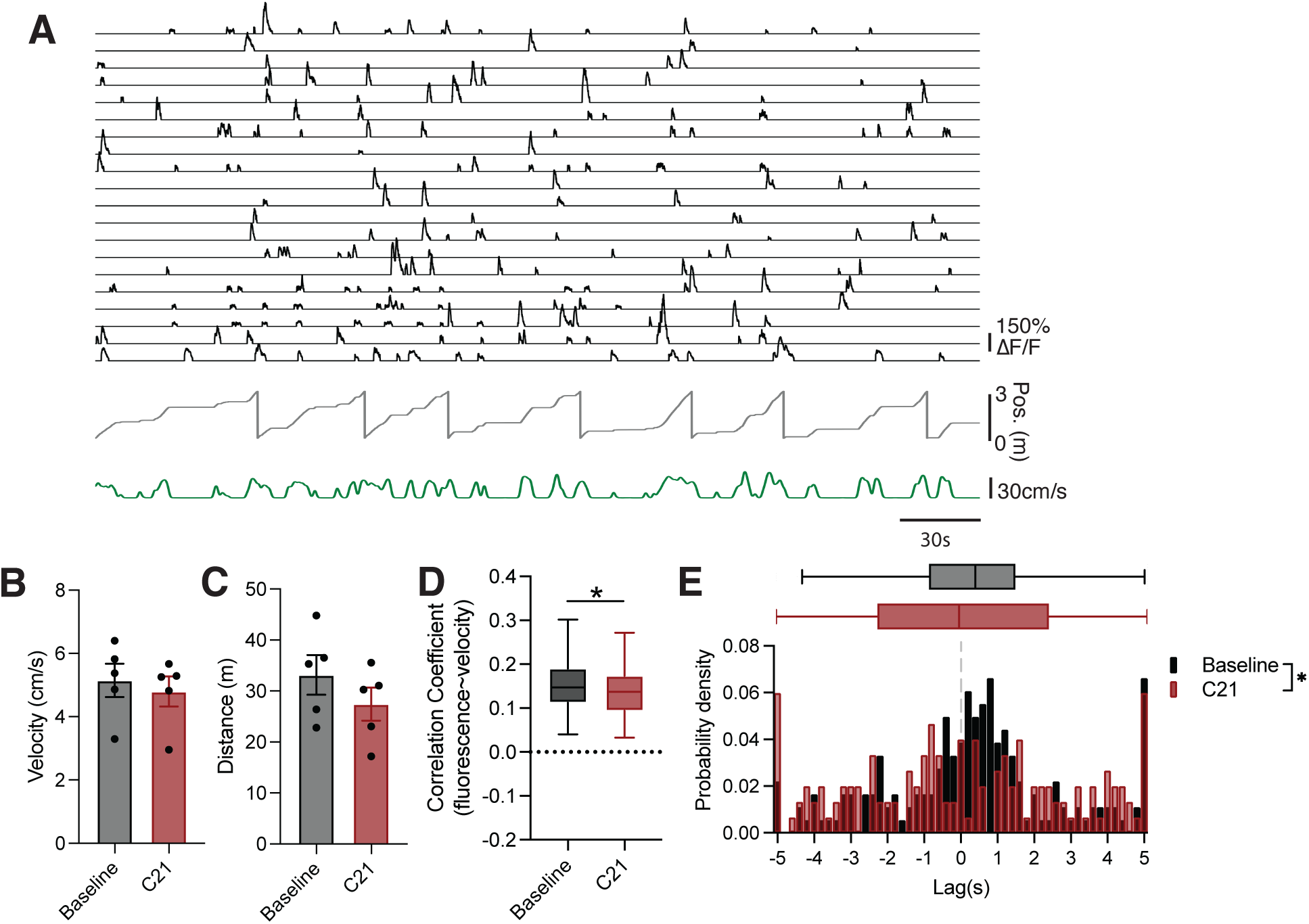
Chemogenetic activation of VGluT3+ interneurons alters coupling of MC activity to velocity. **A)** Example traces of GCaMP fluorescence traces from MCs (black, top), animal’s position in VR (gray, middle) and running velocity (green, bottom). **B)** Bar graph showing animal’s average running velocity during baseline and C21 sessions (n=5 mice; mean± s.e.m.). **C)** Bar graph showing total distance travelled during 12-minute imaging session under baseline and C21 conditions, mean± s.e.m.. **D)** Graph showing the activity-velocity correlation coefficient for MCs during baseline and C21 sessions (baseline, n=181 cells, 5 mice; C21, n = 150 cells, 5 mice; *p<0.05 Mann-Whitney U test). **E)** Histogram showing distribution of activity-velocity correlation lags for MCs during baseline and C21 sessions (baseline, n=181 cells, 5 mice; C21, n = 150 cells, 5 mice; *p<0.05 Kolmogorov-Smirnov test on distributions).

**Supplemental figure 5.**
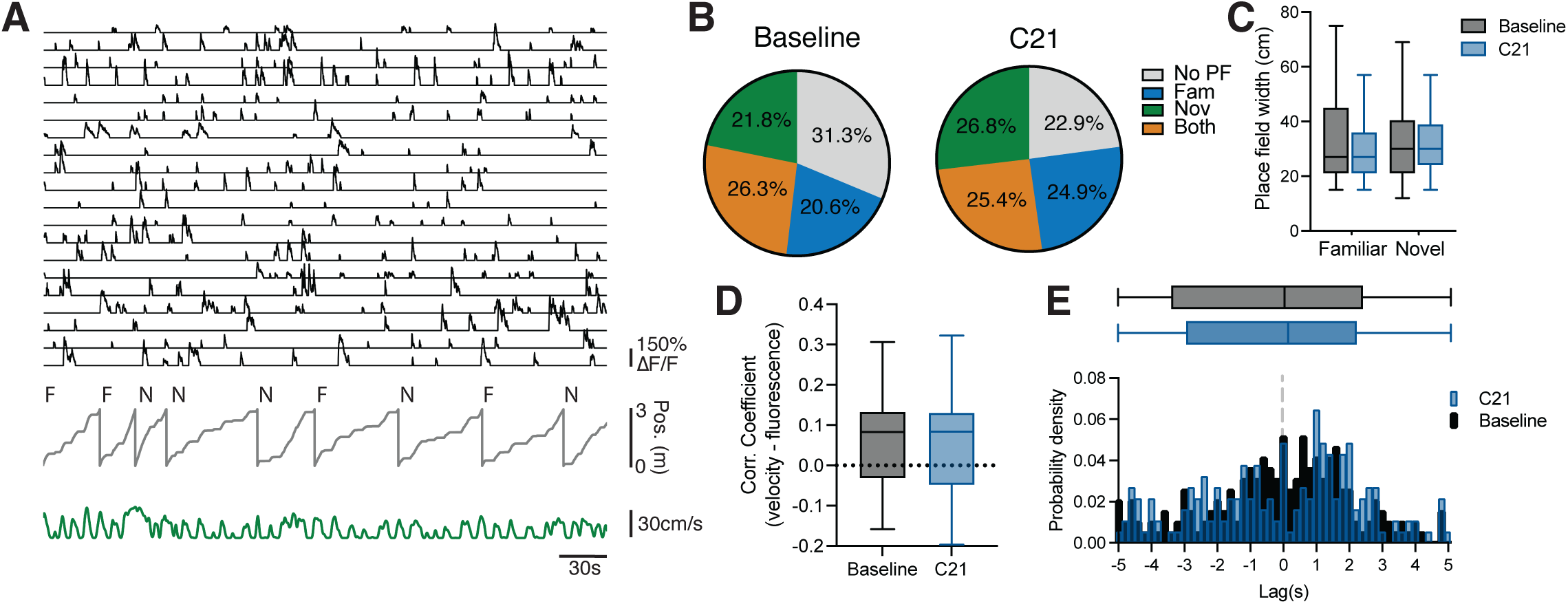
Edect of inhibition of VGluT3+ interneurons on spatial tuning and activity-velocity correlations. **A)** Example traces of GCaMP fluorescence traces from MCs (black, top), animal’s position in VR (gray, middle) and running velocity (green, bottom). Letters above position indicate environment type (Familiar - F or Novel - N) for each lap. **B)** Pie chart showing proportion of spatially tuned MCs in F, N or both environments during baseline and C21 conditions (baseline, n=243 cells, 3 mice; C21, n=201 cells, 3 mice). **C)** Average place field width for spatially tuned MCs during baseline and C21 conditions (baseline, n=243 cells, 3 mice; C21, n=201 cells, 3 mice). **D)** Graph showing activity-velocity correlation coefficient for MCs during baseline and C21 sessions. **E)** Histogram showing distribution of activity-velocity correlation lag during baseline and C21 sessions (baseline, n=243 cells, 3 mice; C21, n=201 cells, 3 mice)

